# Drugs Form Ternary Complexes with Human Liver Fatty Acid Binding Protein (FABP1) and FABP1 Binding Alters Drug Metabolism

**DOI:** 10.1101/2024.01.17.576032

**Authors:** King Clyde B. Yabut, Alice Martynova, Abhinav Nath, Benjamin P. Zercher, Matthew F. Bush, Nina Isoherranen

## Abstract

Liver fatty acid binding protein (FABP1) binds diverse endogenous lipids and is highly expressed in the human liver. Binding to FABP1 alters the metabolism and homeostasis of endogenous lipids in the liver. Drugs have also been shown to bind to rat FABP1, but limited data is available for human FABP1 (hFABP1). FABP1 has a large binding pocket and multiple fatty acids can bind to FABP1 simultaneously. We hypothesized that drug binding to hFABP1 results in formation of ternary complexes and that FABP1 binding alters drug metabolism. To test these hypotheses native protein mass spectrometry (MS) and fluorescent 11-(dansylamino)undecanoic acid (DAUDA) displacement assays were used to characterize drug binding to hFABP1 and diclofenac oxidation by cytochrome P450 2C9 (CYP2C9) was studied in the presence and absence of hFABP1. DAUDA binding to hFABP1 involved high (K_d,1_=0.2 µM) and low affinity (K_d,2_ >10 µM) binding sites. Nine drugs bound to hFABP1 with K_d_ values ranging from 1 to 20 µM. None of the tested drugs completely displaced DAUDA from hFABP1 and fluorescence spectra showed evidence of ternary complex formation. Formation of DAUDA-diclofenac-hFABP1 ternary complex was verified with native MS. Docking placed diclofenac in the portal region of FABP1 with DAUDA in the binding cavity. Presence of hFABP1 decreased the k_cat_ and K_m,u_ of diclofenac with CYP2C9 by ∼50% suggesting that hFABP1 binding in the liver will alter drug metabolism and clearance. Together, these results suggest that drugs form ternary complexes with hFABP1 and that hFABP1 interacts with CYP2C9.

**Significance statement:** Many commonly prescribed drugs bind FABP1 forming ternary complexes with FABP1 and the fluorescent fatty acid DAUDA. This suggests that in the human liver drugs will bind to apo-FABP1 and fatty acid bound FABP1. The high expression of FABP1 in the liver and binding of drugs to FABP1 will alter rates of drug metabolism in the liver.

## Introduction

Fatty acid binding proteins (FABPs) are intracellular lipid binding proteins broadly expressed in tissues (Smathers and Petersen, 2011; Yabut and Isoherranen, 2023). They bind essential endogenous lipids such as fatty acids, bile acids, cholesterol and eicosanoids (Smathers and Petersen, 2011; Yabut and Isoherranen, 2023). FABPs are critical for lipid homeostasis and signaling in variety of tissues through regulation of the uptake, metabolism and cellular trafficking of their ligands (Smathers and Petersen, 2011; Yabut and Isoherranen, 2023). FABPs can also play a role in the pharmacological effects of drugs that bind FABPs. FABPs impact nuclear receptor activation by hypolipidemic drugs (Hughes *et al*., 2015) and alter behavior and cognition associated with cannabinoid signaling (Elmes *et al*., 2019; Penman *et al*., 2023). Changes in FABP expression in the intestines and brain result in altered tissue uptake and disposition of drugs (Trevaskis *et al*., 2011; Penman *et al*., 2023). Yet, little is known about how drug binding to FABPs in the liver alters drug metabolism and liver uptake.

FABP1 is the predominant FABP in the liver. It constitutes 7-10% of all cytosolic protein in the human liver (0.7-1 mM) (Wang *et al*., 2015) and accounts for ∼80% of long chain fatty acid (LCFA) binding in the liver cytosol (Schroeder *et al*., 2016). Despite the extensive characterization of binding of endogenous ligands to FABP1, drug binding to human FABP1 (hFABP1) is not well characterized. Nonsteroidal anti-inflammatory drugs (NSAIDs), fibrates, benzodiazepines, glitazones, β-blockers, steroids and psychoactive cannabinoids bind to rat FABP1 (rFABP1) (Chuang *et al*., 2008; Huang *et al*., 2018). However, rFABP1 and hFABP1 have distinct structural and biochemical differences that likely result in different ligand binding specificities and affinities. rFABP1 and hFABP1 share only 83% amino acid identity with 10% of the sequence being nonconservative amino acid replacements (Schroeder *et al*., 2016). hFABP1 is less alpha helical, has a larger binding cavity, higher thermal stability, and different binding affinities with long chain fatty acid (LCFA) than rFABP1. For drugs, fenofibrate and fenofibric acid bound to hFABP1 with 7 to 23-fold greater binding affinity when compared to rFABP1 (Martin *et al*., 2013). Similarly, some cannabinoids bound to hFABP1 (Elmes *et al*., 2019) but no binding was detected to rFABP1 (Huang *et al*., 2018). Hence, data of drug binding to rFABP1 may not translate to hFABP1 and thorough understanding of general drug binding kinetics with hFABP1 is needed.

Fluorescence displacement assays are widely used to identify FABP ligands and characterize ligand binding to FABPs (Thumser and Wilton, 1994; Velkov *et al*., 2007; Chuang *et al*., 2008; Zhou *et al*., 2019; Yabut and Isoherranen, 2023). However, FABP1 has a large binding cavity and multiple endogenous ligands have been shown to bind FABP1 simultaneously (Santambrogio *et al*., 2013; Favretto *et al*., 2015). This suggests that in fluorescence displacement assays drug ligands may only partially displace the fluorescent ligand leading to a loss of assay sensitivity and potential confounding effects in assessment of ligand binding affinity. With FABP2 such effects were shown with ketorolac (Patil *et al*., 2014). Ketorolac did not displace the fluorescent probe 8-anilino-1-naphthalenesulfonic acid (ANS) from FABP2 and NMR analysis suggested that ketorolac and ANS bind simultaneously to FABP2 (Patil *et al*., 2014). NMR studies have also suggested that drugs form ternary complexes with rFABP1 (Chuang *et al*., 2008). Based on these findings we hypothesized that drug ligands form ternary complexes with hFABP1 either with a fluorescent probe, or with two drug molecules binding simultaneously. To test this hypothesis, we developed a DAUDA displacement assay with singular value decomposition (SVD) analysis in conjunction with native mass spectrometry to characterize ligand binding to hFABP1.

FABP1 has profound effects on the metabolism of endogenous ligands in the liver. FABP1 knockout mice have decreased hepatic fatty acid β-oxidation, decreased triglyceride formation, decreased [^3^H]oleate incorporation into cellular triglycerides and diacylglycerol, and altered hepatic lipid profiles (Martin *et al*., 2003, 2005; Newberry *et al*., 2003; Storch and Corsico, 2008). In perfused rat livers, higher FABP1 expression resulted in higher clearance of palmitate (Hung *et al*., 2003). Notably, FABP1 also interacts directly with carnitine palmitoyl transferase I (CPTI) facilitating LCFA-CoA metabolism (Hostetler *et al*., 2011). Consistent with a role of FABP1 facilitating metabolism, FABP1-knockout mice had decreased rates of Δ9-tetrahydrocannabinol (THC) metabolism (Elmes *et al*., 2019). Based on these data we hypothesized that the metabolism of drugs that bind to hFABP1 is altered in the presence of FABP1 binding. This hypothesis was tested using diclofenac metabolism by recombinant CYP2C9 as a model reaction.

## Materials and Methods

### Chemicals and Reagents

Kanamycin, Trizma base (Tris), sodium chloride, sodium phosphate, potassium phosphate, protease inhibitor tablets, benzonase, thrombin, Coomassie Brilliant Blue R, 11-(Dansylamino)undecanoic acid (DAUDA), arachidonic acid, diazepam, diclofenac, fluoxetine, racemic flurbiprofen, gemfibrozil, ibuprofen, sulfaphenazole and tolbutamide were purchased from Millipore-Sigma (St. Louis, MO). (R)- and (S)-flurbiprofen were purchased from Cayman Chemical (Ann Arbor, MI). Pioglitazone was purchased from Altan Biochemicals. Tryptone, yeast extract, IPTG, PMSF, imidazole, BCA protein assay and low melt agarose were from Thermo Fisher Scientific (Waltham, MA). SeaKem agarose was purchased from Lonza (Basel, Switzerland). Lipidex-5000 slurry in methanol was purchased from Perkin Elmer Inc (Waltham, MA, USA). Mini-PROTEAN TGX protein gels were purchased from Bio-Rad (Hercules, CA). HindIII and NdeI restriction enzymes were purchased from New England BioLabs (Ipswich, MA). Lyophilized ribonuclease A was purchased from Sigma Aldrich (St. Louis, MO). Ultrapure ammonium acetate salt was purchased from VWR Scientific (San Francisco, CA) and tuning mix for ESI-time-of-flight mass spectrometry was purchased from Agilent (Santa Clara, CA). 4’OH-diclofenac and 4’OH-5-chloro-diclofenac were a gift from Dr. Allan Rettie (Department of Medicinal Chemistry, University of Washington).

### Expression, Purification and Delipidation of Human FABP1

Hexa-histidine tagged human FABP1 was expressed in Rosetta 2 *E. coli* (Novagen, Madison, WI) and FABP1 purification and delipidation were optimized and conducted as described in detail in Supplemental Materials. In brief, the his-tagged FABP1 was purified using HisTrap HP affinity column (GE Healthcare, Chicago, IL), the tag was cleaved by thrombin and the cleaved protein was purified and buffer exchanged by gel filtration into 10 mM potassium phosphate pH 7.4, 150 mM KCl. The FABP1 was delipidated using butanol and Lipidex-5000 (Perkin Elmer Inc., Waltham, MA, USA). FABP1 was stored on ice and the concentration was quantified via bicinchoninic acid (BCA) (Pierce, Waltham, MA) assay prior to adding 0.5 mM DTT. The purified protein was characterized using native mass spectrometry.

### Fluorescence Assay for DAUDA Binding to FABP1

All fluorescence spectra were collected using a Cary Eclipse fluorescence spectrophotometer (Agilent, Santa Clara, CA). Spectra were collected in 2 mL of assay buffer (100 mM potassium phosphate, pH 7.4) in a 4 mL clear quartz cuvette at 21°C. The final concentration of organic solvent was kept <1.6%. DAUDA binding to FABP1 was monitored via the enhancement of DAUDA fluorescence due to binding to FABP1 using an excitation wavelength of 335 nm and by monitoring emission from 400-700 nm. All experiments were repeated on at least three separate days and with at least two independent batches of purified protein. Detailed description of the fluorescence spectroscopy is included in Supplemental Materials.

The equilibrium binding affinity constant (K_d_) for DAUDA with FABP1 was determined using reverse and forward fluorescence titrations. A range of concentrations of FABP1 and DAUDA were initially tested to optimize experimental conditions based on detector sensitivity and ligand binding/depletion. Reverse titrations were then performed with a constant concentration of DAUDA (0.05 µM) and increasing concentrations of FABP1. Forward titrations were performed with a constant concentration of FABP1 (0.3 µM) and increasing DAUDA concentrations. The emission spectrum of DAUDA in solution overlaps with that of DAUDA-FABP1 (Figure S6A) preventing direct measurement of DAUDA-FABP1 fluorescence in titration experiments. Singular value decomposition (SVD) was therefore used to deconvolute fluorescence titration spectra. SVD can be used to analyze spectral data and quantify contributions from spectrally distinct species measured over the course of a titration (Hendler and Shrager, 1994; Nath *et al*., 2008). SVD yields a set of singular values that reveal how many spectrally distinct species contribute to a titration: if there are *n* species that make independent contributions, there will be *n* singular values that are significantly greater than 0 (all subsequent singular values will be close to 0, and simply reflect noise in the data). Spectral deconvolution requires selection of basis spectra corresponding to the individual species that contribute to the observed fluorescence signal. The details of the SVD analysis including construction of basis spectra are provided in Supplemental Materials.

Basis spectra of DAUDA in solution and DAUDA-FABP1 complex were used for deconvolution of titration spectra and determination of the specific fluorescence of DAUDA-FABP1 complex. Binding isotherms were generated by plotting the specific fluorescence of DAUDA-FABP1 against total DAUDA concentration. The high affinity equilibrium dissociation constant (K_d,1_) for DAUDA binding to FABP1 was determined by fitting the reverse titration data to the following tight binding equation:

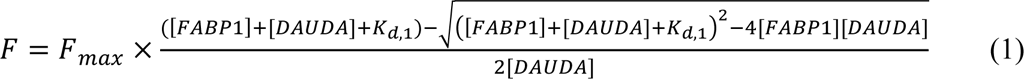

In equation 1, F is the fluorescence of DAUDA-FABP1 determined by SVD, F_max_ is the maximum fluorescence of DAUDA-FABP1, and [FABP1] and [DAUDA] are the total concentrations of FABP1 and DAUDA used in reverse titration experiments, respectively. Results from the quadratic equation fit were verified by fitting a numerical model of bimolecular association to the data using COPASI (Hoops *et al*., 2006). The low affinity binding site (K_d,2_) was determined by fitting a two-site sequential binding model to the forward titration data in COPASI using K_d,1_ from the reverse titration fit.

For the reverse titration, the K_d,1_ of DAUDA with FABP1 was determined by fitting the following single binding site model to the steady-state fluorescence data in COPASI:

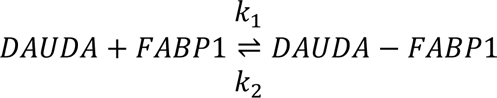

A scaling factor (*Scale*) was used to relate the observed fluorescence *F* of to the concentration of the DAUDA-FABP1 yielded by the kinetic model:

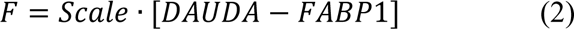

In equation 2, [DAUDA-FABP1] is the equilibrium concentration of DAUDA-FABP1 throughout the course of the titration. The initial concentration of DAUDA was constrained to 0.05 µM and the association rate constant (k_1_) was constrained to 1 µM^-1^ s^-1^ The dissociation rate constant (k_2_) and *Scale* were fitted parameters. The lower and upper bound values for k_2_ were 0.1 and 10, respectively, and the lower and upper bound values for *Scale* were 1,000-1,000,000, respectively. K_d,1_ was determined as the ratio of k_2_/k_1._ The results of the fit were insensitive to the start values for k_2_ and *Scale*.

For the forward titration, the following two-site sequential binding model was fit to the fluorescence data using COPASI:

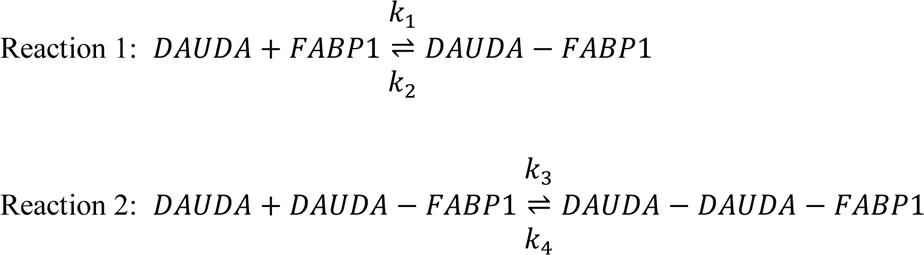

The total fluorescence (*F_tot_*) observed from singly and doubly bound DAUDA-FABP1 complexes was defined by the following equation:

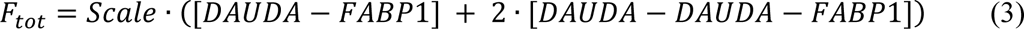

Here, *Scale* is a scaling factor for the fluorescence of DAUDA-FABP1 complexes, with the assumption that the doubly-bound complex fluorescence is twice as intense as the singly-bound. [DAUDA-FABP1] and [DAUDA-DAUDA-FABP1] are the equilibrium concentrations of singly and doubly bound DAUDA-FABP1 complexes, respectively. For model fitting, the initial concentration of FABP1 was constrained to 0.3 µM. The association and dissociation rate constants (k_1_ and k_2_, respectively) for reaction 1 were fixed to 1 µM^-1^ s^-1^ and 0.2 s^-1^ respectively, to be consistent with the single site binding affinity determined from reverse titrations. The association rate constant for reaction 2 (k_3_) was fixed to 1 µM^-1^ s^-1^ and the dissociation rate constant (k_4_) was a fitted parameter along with *Scale*. The lower and upper bounds for k_4_ were 0.1 and 10, respectively, and the lower and upper bound values for *Scale* were 1,000 and 1,000,000, respectively. The binding affinity for the second reaction (K_d,2_) was determined as the ratio of k_4_/k_3_ and results of the fit were insensitive to the start values for k_4_ and *Scale*. DAUDA binding affinities are reported from a single fit to data combined from replicate experiments performed on separate days and with FABP1 from different purifications.

### DAUDA displacement assay for hFABP1 binding

Arachidonic acid (AA) was first used as a model ligand to develop a method to measure ligand binding to FABP1 via DAUDA displacement. AA in methanol was titrated into a solution of FABP1 (0.3 µM) prebound with DAUDA (0.5 µM). AA was confirmed to have no background fluorescence in buffer or in the presence of 0.3 µM FABP1. The concentrations of DAUDA-FABP1 complex in AA titration experiments were determined by SVD analysis using basis spectrum for DAUDA in buffer and DAUDA-FABP1 as described in Supplemental Materials. To determine the apparent binding affinity of AA (K_d_) with hFABP1, a competitive displacement model (Figure S9A) was fit to the SVD data for AA using COPASI. The competitive displacement model was based on the two-site sequential binding model described above for DAUDA with the addition of a third reaction:

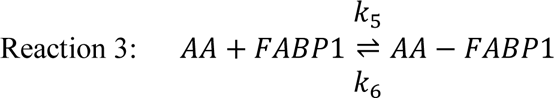

The SVD fluorescence titration data for AA was normalized to the fluorescence of DAUDA bound with FABP1 in the absence of AA. *F_tot_* was defined as in equation 3, and *Scale* was fixed to 529 based on the initial concentrations of DAUDA-FABP1 and DAUDA-DAUDA-FABP1 in displacement assays (Supplementary Materials). k_5_ and k_6_ are the association and dissociation rate constants for AA binding with FABP1, respectively. k_5_ was set to 1 µM^-1^ s^-1^ and k_6_ was fitted with lower and upper bounds of 0.001 and 1000, respectively. The fitted value for k_6_ was independent of the initial value used. The K_d_ from the ratio of k_6_/k_5_ is reported as the mean ± standard deviation from three replicate experiments performed on separate days with at least two different purifications from a single expression of hFABP1.

Drug binding to hFABP1 was screened using simple DAUDA displacement. Diazepam, diclofenac, fluoxetine, racemic flurbiprofen, gemfibrozil, racemic ibuprofen, pioglitazone, sulfaphenazole and tolbutamide solutions were prepared in methanol and added at 30 μM to FABP1 (0.3 µM) prebound with DAUDA (0.5 µM). All drugs were confirmed to have no appreciable background fluorescence at 30 µM in buffer or in the presence of 0.3 µM FABP1. For ligands that reduced the DAUDA-FABP1 fluorescence >15% in initial screening, titrations were performed with a range of ligand concentrations added to FABP1 (0.3 µM) prebound with DAUDA (0.5 µM). Fluorescence measurements and SVD analysis were performed as described for AA above. (R)- and (S)-flurbiprofen were used in titrations instead of racemic flurbiprofen.

Because residual DAUDA-FABP fluorescence was observed even at saturating concentrations of many ligands, a ternary complex binding model (Figure S9B) was used in COPASI to determine the binding affinity of drugs (K_d_) with hFABP1. The ternary binding model was based on the two-site sequential binding model described above for DAUDA (Reactions 1 and 2) with the addition of a third reaction:

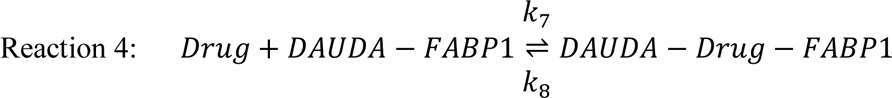

cThis is the simplest model that adequately fits the data but relies on the assumption that drug binding does not alter the affinity of DAUDA for either its high- or low-affinity binding sites. The SVD data from drug titrations was normalized to the fluorescence of DAUDA bound with FABP1 in the absence of drug. The total fluorescence *F_tot_* observed from singly and doubly bound DAUDA-FABP1 complexes and DAUDA-Drug-FABP1 ternary complexes was defined by the following equation:

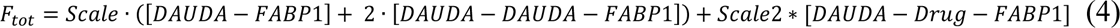

*Scale2* is a scaling factor for the fluorescence of the DAUDA-drug-FABP1 ternary complex. *Scale* was fixed to 529 as described above for AA. k_7_ and k_8_ are the association and dissociation rate constants for drug binding with FABP1, respectively. k_7_ was set to 1 µM^-1^ s^-1^ k_8_ and *Scale2* were allowed to vary. The lower and upper bounds for k_8_ were set to 0.001 and 1000, respectively, and the initial value was fixed to the EC_50_ determined as described below. The lower and upper bounds for *Scale2* were set to 0 and 1000, respectively, and an initial value of 500 was used. The K_d_ values for drug ligands were determined from the ratio of k_8_/k_7_ and are reported as means ± standard deviation from three replicate experiments done on separate days with at least two different purifications of FABP1.

EC_50_ values were also determined as an alternative measurement of drug binding affinity. To determine the concentration of ligand at half maximal displacement of DAUDA (EC_50_ value), the % fluorescence remaining for DAUDA-FABP1 as determined by SVD analysis was plotted as a function of ligand concentration. Equation 5 was fit to the data in GraphPad Prism 10.

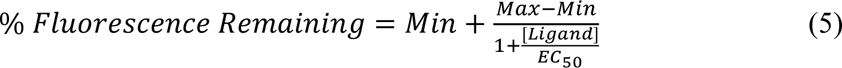

Here, [Ligand] is the concentration of AA or the test drug and Min and Max are the minimum and maximum values for % fluorescence remaining, respectively. Min values were constrained to be > 0 and Max values were fixed to 100. The EC_50_ values are reported as a mean ± standard deviation from replicate experiments done on three separate days with FABP1 from different purifications. The residual fluorescence remaining (F_res_) at saturating drug concentrations was taken as *Min* from equation 5.

### Native mass spectrometry methods for characterization of DAUDA and diclofenac binding to FABP1

Non-delipidated and delipidated FABP1 samples were prepared for native mass spectrometry using Micro Bio-Spin Size-Exclusion Spin Columns (Bio-Rad, Hercules, CA) that had been equilibrated with four washes of 1M ammonium acetate adjusted to pH 7. FABP1 aliquots were diluted in 1M ammonium acetate to 50 µL prior to loading onto the equilibrated column. FABP1 was eluted from the column by centrifugation at 1,000 g for four and a half minutes. Assuming 100% recovery from the column and negligible protein loss due to adsorption, the volume of recovered FABP1 solution was then measured and diluted with additional 1 M ammonium acetate and DTT to reach a final concentration of 10 µM FABP1 and 10 mM DTT. FABP1 and FABP1-ligand complex ions were generated using nano-electrospray ionization (Davidson *et al*., 2017). Mass spectrometry analysis was performed on a Q-Tof Premier Mass Spectrometer (Waters Corp., Wilmslow, UK). Ion source and transfer conditions were optimized to minimize ion activation.

To characterize ligand binding, DAUDA and diclofenac dissolved in methanol were pipetted directly into 1 M ammonium acetate solution of FABP1 (10 µM) to achieve the desired stoichiometric molar ratios of ligand and protein while keeping the final concentration of methanol below 5% by volume. After addition of ligands, samples were allowed to equilibrate at 4 °C overnight. Approximately 1-3 µL of protein solution was loaded into glass emitters that were made in-house using borosilicate capillaries and a micropipette puller (Sutter Instruments Model P-97, Novato, CA). A platinum wire was inserted into the solution and approximately 0.5-1 kV was applied to the wire to generate ions. Native mass spectra were acquired with a 35 V bias between the sampling and extraction cones in the ion source, which was operated at room temperature. A 3-5 V bias was applied between the quadrupole mass filter and the entrance to the collision cell, which were the least activating setting that allowed for sufficient ion transmission. External calibration of the mass spectra was performed using nano-electrospray generated ions from a commercially available calibration standard (Agilent, Santa Clara, CA). Native mass spectra were manually processed using MassLynx (Waters Corp., Wilmslow, UK) and software written for this project.

In native mass spectrometry analyses, nonspecific ligand binding can occur during the electrospray process due to concentration effects (Kitova *et al*., 2012). One well documented method to verify the specificity of an interaction between a protein (P) and ligand (L) is the reference protein experiment (Sun *et al*., 2006). In this experiment, an additional protein (P_ref_) is added to the sample solution that is not expected to interact with L. Any observed peaks corresponding to P_ref_+L are attributed to nonspecific interactions that are an artifact of the electrospray process and can be subtracted from P+L peaks (Kitova *et al*., 2012). Using this strategy, the native mass spectrum of FABP1 with DAUDA was generated with the addition of ribonuclease A (data not shown) as reference protein. These experiments were performed by adding 10 µL of 20 µM ribonuclease A in 1 M ammonium acetate during the final dilution of FABP1 to 10 µM. The total volumes of the FABP1 solutions with and without ribonuclease A were the same, ensuring consistent relative concentrations of FABP1 and ligands. DAUDA and diclofenac were added to FABP1/ribonuclease A solutions and allowed to equilibrate at 4°C overnight prior to native mass spectrometry analysis. There was no evidence for DAUDA association with ribonuclease A, while both singly and doubly bound DAUDA with FABP1 peaks were present. These results indicate that DAUDA-associated peaks in the native mass spectra result from specific condensed-phase interactions.

### Molecular Docking of DAUDA and Drugs to FABP1

DAUDA, diclofenac, (R)-flurbiprofen and (S)-flurbiprofen were docked to a holo-FABP1 solution structure determined by NMR and in complex with two oleic acid molecules (PDB 2LKK, chain A.1) (Cai *et al*., 2012). Docking was performed with AutoDock4 using AutoDock Tools (1.5.7) (Rizvi *et al*., 2013). Protonated 3D structures of the drugs and endogenous ligands were downloaded from PubChem. Polar hydrogens and Kollman charges were added to FABP1 using AutoDock Tools (1.5.7). Ligand torsions were automatically selected with AutoDock Tools and verified according to the 3D ligand structure, and the ligand aromaticity criterion was set to 7.5. Grid parameter files were prepared using a grid box size of 80×100×90 (X, Y, Z) centered on FABP1 to encompass the entire β-barrel binding domain of FABP1. Docking parameter files were prepared using a rigid structure of FABP1. The Genetic Algorithm (GA) was used with default settings with the number of GA runs set to 50. The docking parameters were set as the default and the docking parameter file was output as LamarkianGA (4.2). For docking studies with two ligands, a single ligand was first docked to the holo-FABP1 structure, and the top scoring pose (lowest ΔG_binding_) was used as the holo-FABP1 structure for additional docking studies with a second ligand. Docking poses were visualized using ChimeraX 1.1 (University of California, San Francisco) (Pettersen *et al*., 2004).

### Kinetics of 4’OH-diclofenac Formation by CYP2C9 in the Presence and Absence of FABP1

The kinetics of 4’OH-diclofenac formation by CYP2C9 in the presence and absence of hFABP1 were determined in CYP2C9 Supersomes under conditions of protein and time linearity. CYP2C9 Supersomes (1 nM CYP2C9, 0.0015 mg total microsomal protein/mL) were preincubated with 8 different concentrations of diclofenac ranging from 0.4 to 20 µM for 5 min at 37°C in 180 µL incubation buffer (100 mM potassium phosphate, pH 7.4) in a 96-well plate. Reactions were initiated with 1 mM NADPH (final concentration) to a final volume of 200 µL and quenched after 10 min by transferring 100 µL of the incubation to 0.6 mL Eppendorf tubes containing three incubation volumes of acetonitrile with 1% formic acid and 17 nM of 4’OH-5-chloro-diclofenac as an internal standard. The incubations in the presence of hFABP1 were conducted in a similar manner as those without hFABP1. For incubations with diclofenac and hFABP1, CYP2C9 was preincubated with diclofenac and 20 µM hFABP1 for all concentrations of diclofenac tested prior to initiation of the catalytic reactions with NADPH.

For incubations in the presence and absence of FABP1, quenched reactions were centrifuged at 18,000 g for 20 minutes at 4°C and 200 µL of supernatant was collected and transferred to glass mass spectrometry (MS) vials for LC-MS/MS analysis. Diclofenac samples were analyzed using an AB Sciex API6500 qTrap mass spectrometer (Concord, ON, Canada) coupled to an Agilent 1290 Infinity II Ultra-High-Performance Liquid Chromatography (UHPLC, Santa Clara, CA). For 4’OH-diclofenac separation, a Synergi Max-RP column (150 x 4.6 mm, 4 μM, Phenomenex, Torrance, CA) was used. A gradient elution at a flow rate of 1 mL/min was used as follows: mobile phase A (water with 0.1% formic acid) was kept at 65% and B (acetonitrile with 0.1% formic acid) at 35% for the first 18 minutes, then B was increased to 80% by 25 minutes, returned to initial conditions by 30 minutes and held at initial conditions for an additional 5 minutes. 4’OH-diclofenac and 4’OH-5-chloro-diclofenac were monitored in positive ion mode with electrospray ionization and the MS parameters used were as follows: IS 4500 V, TEM 400 °C, CUR 35 p.s.i., GS1 62, GS2 62, CAD-low, EP 10 V, DP 60 V, CXP 14 V, and the CE were 22 and 19 V for 4’OH-diclofenac and 4’OH-5-chloro-diclofenac, respectively. The MRM transitions used were 312>266 *m/z* for 4’OH-diclofenac and 346>300 *m/z* for 4’OH-5-chloro-diclofenac.

### Determination of Diclofenac Unbound Fraction in CYP2C9 Incubations

To determine the unbound concentrations of diclofenac in incubations with recombinant CYP2C9, magnetic silica beads (MGSBs, G-Biosciences, St. Louis, MO) were used to separate microsomal protein (Horspool *et al*., 2020) from free diclofenac in solution. Prior to experiments the beads were conditioned and washed with 3 mL (1 mL x 3) of assay buffer (100 mM potassium phosphate, pH 7.4). Initial experiments without microsomal protein present were done to verify that diclofenac and FABP1 did not bind non-specifically to MGSBs. To measure non-specific binding of diclofenac and FABP1 to MGSBs, 1.9 µM diclofenac or 10 µM FABP1 were incubated separately with 100 µL of MGSB in 0.5 mL assay buffer in 1.7 mL Eppendorf tubes for 30 minutes at 37°C. After 30 minutes, 100 µL of the mixture containing MGSBs with diclofenac or FABP1 were collected as the total sample, then the MGSBs were separated from solution using a DynaMag-2 Magnet (Thermo Fisher Scientific, Waltham, MA) and the supernatant was collected. For supernatant containing FABP1, FABP1 was quantified using BCA protein assay. For diclofenac samples, 300 µL of acetonitrile containing 1% formic acid and 1 µM 4’OH-5-chloro-diclofenac internal standard were added to 100 µL of the total and supernatant samples containing diclofenac. The samples were centrifuged at 18,000 g for 20 minutes and the supernatant was transferred to MS vials for analysis. Diclofenac concentrations in the samples were measured using a Synergi Max-RP column (150 x 4.6 mm, 4 µM, Phenomenex, Torrance, CA) coupled to an Agilent 1200 Series High-Performance Liquid Chromatography (HPLC)-UV system (Santa Clara, CA). A flow rate and elution gradient for diclofenac were used as described above. Diclofenac and 4’OH-5-chloro-diclofenac UV absorbances were monitored at 280 nm and integration of the peaks was done in ChemStation B.04.02 (Agilent, Santa Clara, CA).

To determine the free concentrations of diclofenac in incubations with CYP2C9, 100 μL of MGSBs (10.5 x 10^9^ beads) (Horspool *et al*., 2020) were washed 3 times with 1 mL of assay buffer and pre-equilibrated with CYP2C9 Supersomes (1 nM CYP2C9, 0.0015 mg total microsomal protein/mL) on ice for 30 minutes in 0.5 mL of assay buffer in 1.7 mL Eppendorf tubes. Diclofenac was then added to the mixture of MGSBs and CYP2C9 at concentrations corresponding to each of the nominal concentrations used in kinetic experiments. Samples were then incubated for an additional 30 minutes in a shaking water bath at 37°C, then removed from the water bath and cooled at room temperature for 5 minutes. For experiments with FABP1, FABP1 (20 µM) was pre-equilibrated together with CYP2C9 and MGSBs prior to the addition diclofenac. After cooling, 100 μL of the mixture containing the MGSBs, Supersomes, and diclofenac with and without FABP1 were collected as the total drug sample. The MGSBs were then separated from solution using a DynaMag-2 Magnet and 100 μL of supernatant were collected as the free diclofenac or free diclofenac together with FABP1-bound diclofenac sample. 300 μL of acetonitrile containing 1% formic acid and internal standard were added and samples analyzed as described above. Diclofenac concentrations were determined via HPLC-UV as described above. The binding experiments were done as technical duplicates and the data are reported as means ± S.D. from experiments done on three separate days.

Unbound diclofenac concentrations in the absence of FABP1 were determined from supernatant samples and were measured for every diclofenac concentration used in kinetic experiments. The unbound fraction (*f_u_*) was calculated as the ratio of the concentration of drug measured in supernatant (*Cfree)* to the concentration of total drug measured prior to magnetic separation (*Ctotal)*.

The unbound fraction of diclofenac in the presence of FABP1 was directly measured using Pierce Nickel-nitrilotriacetic acid (Ni-NTA) Magnetic Agarose Beads (Thermo Fisher Scientific, Waltham, MA) for all diclofenac concentrations used in kinetic experiments. A solution of 0.5 mL of 6xHis tagged FABP1 (20 μM) in assay buffer was prebound to diclofenac at room temperature for 10 minutes in 1.7 mL Eppendorf tubes. After 10 minutes, the FABP1 and diclofenac solution was added to magnetic Ni-NTA agarose beads (MNABs) that were prewashed 3 times with 1 mL assay buffer. The mixture of FABP1, diclofenac and MNABs were incubated in a shaking incubator at 25 °C for an additional 30 minutes to bind FABP1 to the MNABs. After 30 minutes, 100 μL of the mixture were taken to measure total diclofenac, then the MNABs were separated from solution using a DynaMag-2 Magnet and 100 μL of supernatant was taken to measure free diclofenac concentration in solution. Diclofenac concentrations were determined via HPLC-UV as described above and the fu was calculated. BCA protein assay was used to measure FABP1 in supernatant samples after magnetic separation to verify that FABP1 bound to MNABs.

### Kinetic Analysis of 4’OH-diclofenac Formation by CYP2C9 in the Presence and Absence of FABP1

The Michaelis-Menten model was fit to the 4’OH-diclofenac formation data in GraphPad Prism 10 using nominal and free concentrations of diclofenac to determine the apparent and unbound 4’OH-diclofenac formation kinetics with CYP2C9, respectively. Experiments in the presence and absence of FABP1 were done as matched pairs on the same day with technical duplicates. Km and kcat values are reported as means ± S.D. from experiments done on three separate days. For every replicate experiment, one-tailed Z-test was used to evaluate differences between Km and kcat values for 4’OH-diclofenac formation in the presence and absence of FABP1 and the results were interpreted collectively from the replicate experiments. The upper limit of the 95% confidence interval for apparent Km values in the absence of FABP1 and the lower limit of the 95% confidence interval for the apparent Km values in the presence of FABP1 were used to calculate the standard error. The standard error for unbound Km and kcat values was calculated using the lower limit of the 95% confidence interval for parameter values in the absence of FABP1 and the upper limit of the 95% confidence interval for parameter values in the presence of FABP1. The standard errors were then used to calculate Z-scores to compare Km and kcat values determined with and without FABP1. A p-value < 0.05 was considered significantly different in each comparison and the values were considered different if all three experiments yielded the same conclusion.

## Results

### Expression, Purification, Delipidation and Characterization of Recombinant hFABP1

FABPs are promiscuous proteins that bind diverse ligands including native *E. coli* lipids and molecules present in expression and purification media (Velkov *et al*., 2008; Wang *et al*., 2017). Such ligands may be bound to recombinant purified FABPs as contaminating copurifying molecules (CPMs). These CPMs may alter the binding characteristics of other ligands (Velkov *et al*., 2008) via competition for FABP binding or via allosteric mechanisms. Hence, a method is needed to monitor the presence of CPMs in purified hFABP1. A native protein mass spectrometry method was developed to assess the extent of CPMs bound to hFABP1 at different stages of the purification (Figure 1, Supplemental Figures S3 and S5) and to confirm efficient delipidation of the final purified protein. Native MS is well suited to monitor the presence of CPMs and is more suitable for routine monitoring of delipidation than previously described methods such as protein NMR which requires isotope labeled protein. It is important to note that due to the limitations of the method, the CPM region likely includes also peaks from modifications to the protein which are unrelated to the purification method, peaks related to presence of sodium and potassium adducts, and peak tailing.

**Figure 1:**
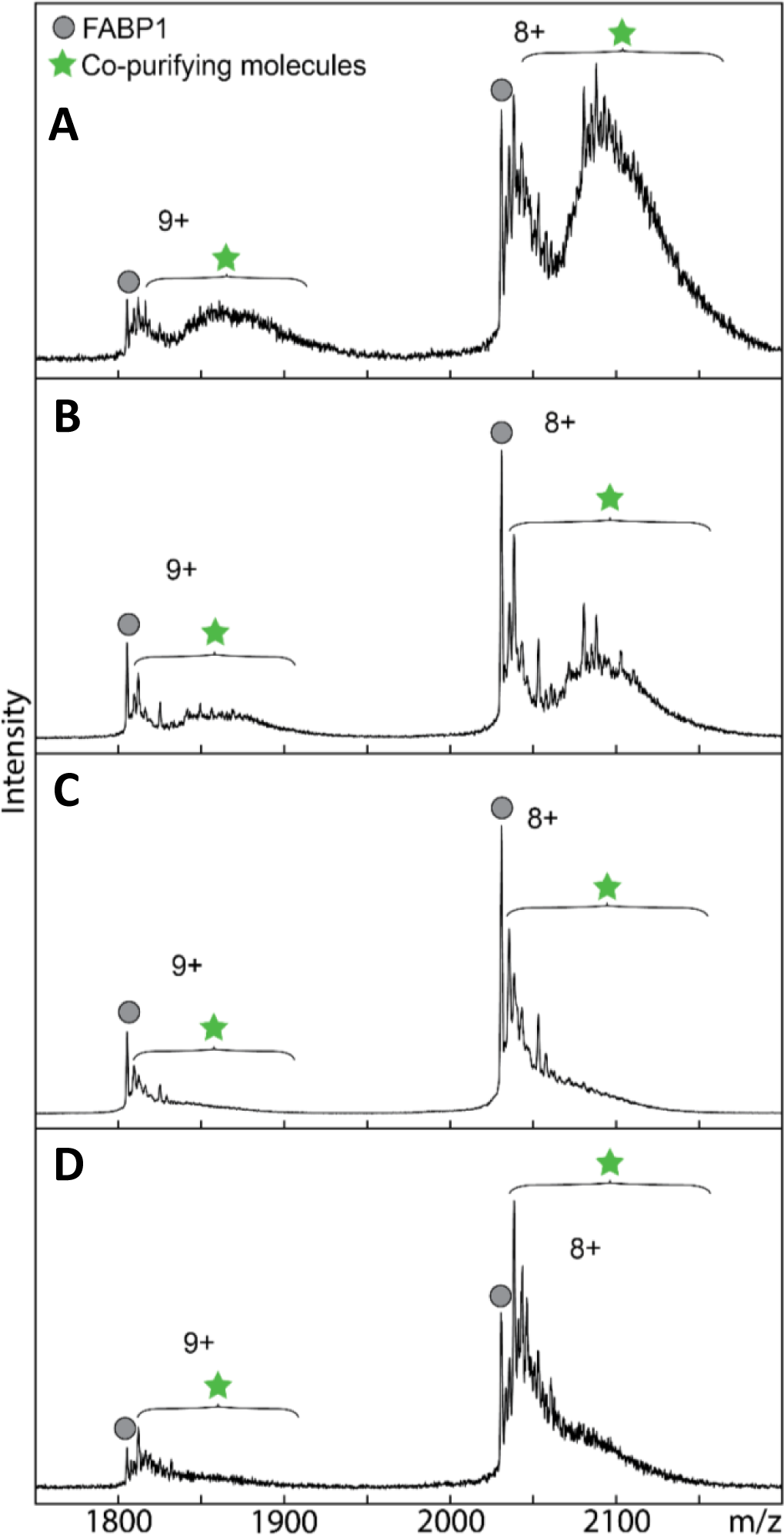
Comparison of FABP1 delipidation methods by native protein mass spectrometry. Native mass spectra of purified his-tagged FABP1 with no delipidation treatment **(A)** and following individual delipidation treatments of Lipidex-5000 **(B),** 1:1 (v/v) butanol **(C)** and 1:3 (v/v) butanol **(D).** Purification and delipidation protocols are described in detail in Supplemental Materials. Mass spectra for FABP1 are shown for the two most abundant charge states (n+). Circle markers denote apo-his-tagged FABP1 and star markers designate the m/z region where co-purifying molecules are observed. The calculated intact mass of his-tagged apo-FABP1 (16,372 Da) aligns with the predicted mass from the amino acid sequences.

In the preliminary experiments, majority of the hFABP1 was observed bound with CPMs after nickel purification and before gel filtration and delipidation treatments (Figure 1A). Lipidex-5000 and butanol extraction have been reported to efficiently delipidate FABP1 (Velkov *et al*., 2008; Wang *et al*., 2017; Lai *et al*., 2020). Various levels of CPMs were removed with individual treatments with Lipidex-5000, 1:1 (v/v) butanol and 1:3 (v/v) butanol based on the native MS analysis (Figure 1). Surprisingly, none of the individual treatments achieved complete delipidation of hFABP1. To accomplish complete removal of the CPMs, a combination of treatments with butanol and Lipidex-5000 were optimized (Supplemental Figure S5). The best efficiency of delipidation was achieved when hFABP1 was treated 3 times with 1:1 butanol followed by a 30-minute incubation with Lipidex-5000 (Supplemental Figure S5D). The final purification protocol is outlined in Figure 2 and the efficiency of the delipidation is shown for the final purified hFABP1.

**Figure 2:**
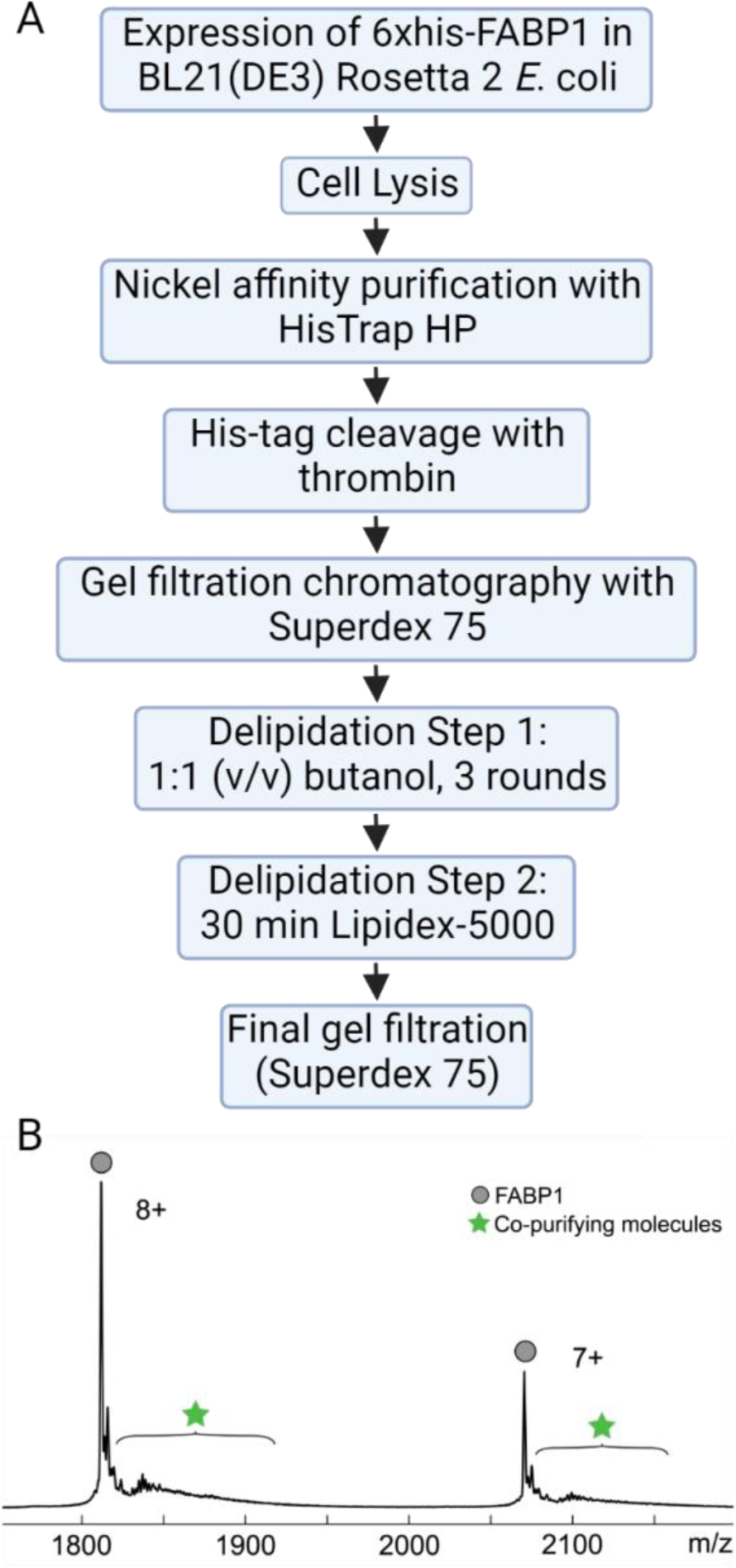
Purification protocol and mass spectrum for recombinant human FABP1. (**A**) Flow chart of final purification protocol for hFABP1 including an optimized delipidation method. (**B**) Mass spectrum of cleaved hFABP1 after final gel filtration step is shown for the two most abundant charge states (n+). Circle markers represent *m/z* spectral peaks for apo-hFABP1 and the star markers indicate the region in the spectra where copurifying molecules are observed or expected to be observed. The calculated intact mass of cleaved apo-hFABP1 (14,489 Da) aligns with the predicted mass from the amino acid sequence. Flow chart in panel A created with BioRender.com.

### Characterization of DAUDA Binding to FABP1

The fluorescence emission spectrum of free DAUDA in solution overlapped with the spectrum of DAUDA bound to hFABP1 (Supplemental Figure S6). The emission peak of DAUDA-FABP1 was observed at 509 nm. Free DAUDA in solution contributes to the total fluorescence signal observed at this wavelength. This fluorescence overlap can confound titration experiments in which the fraction of total DAUDA that is free in solution changes with DAUDA concentration. Hence, singular value decomposition (SVD) analysis was used to distinguish the fluorescence of DAUDA-FABP1 from free DAUDA in solution in titration experiments (Figure 3).

**Figure 3:**
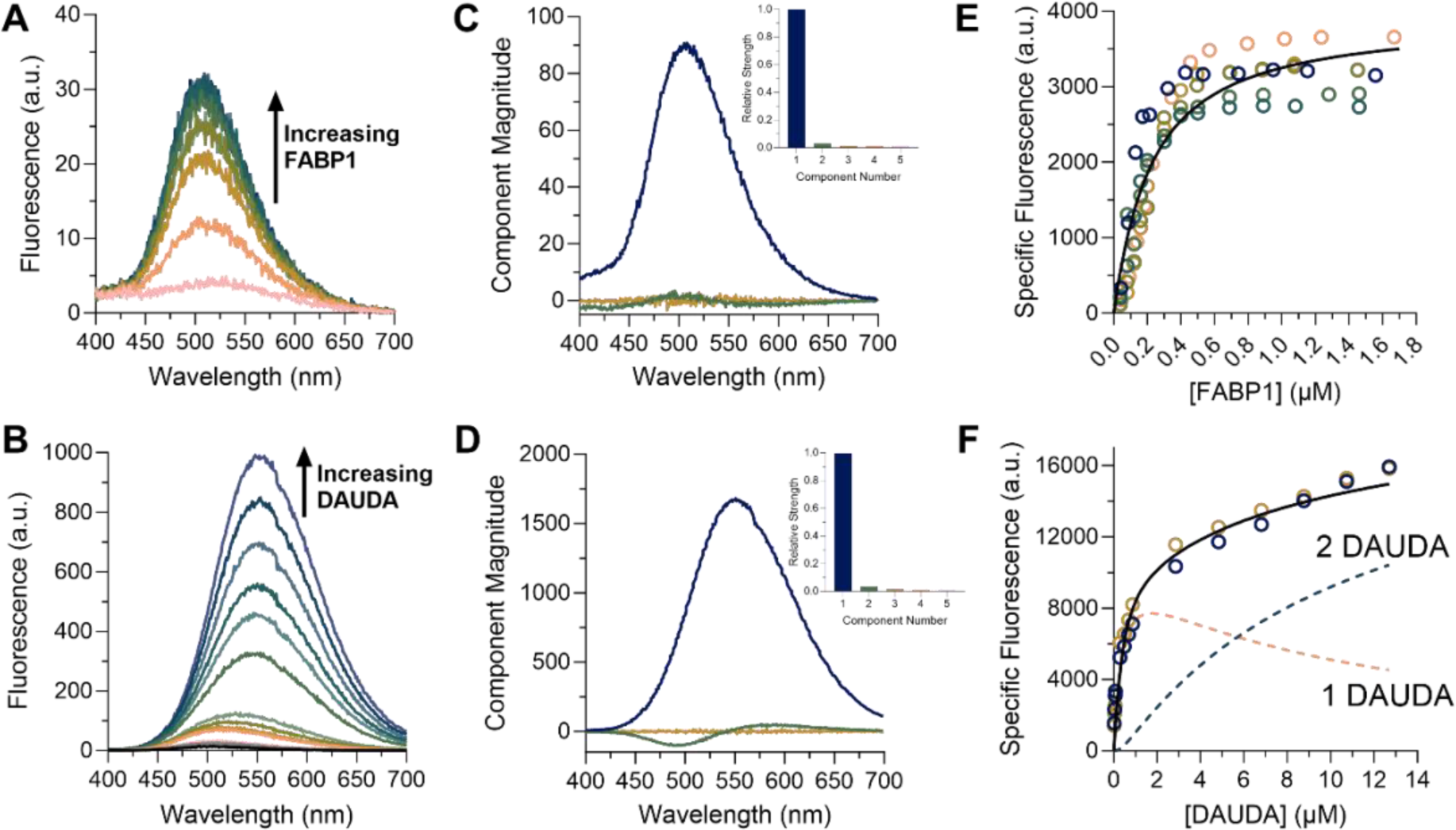
Characterization of DAUDA binding to FABP1 by fluorescence spectroscopy. Fluorescence emission spectra of **(A)** reverse and **(B)** forward titrations. Reverse titrations were done with constant DAUDA (0.05 µM) and increasing FABP1 (0.04-1.7 µM). Forward titrations were done with constant FABP1 (0.3 µM) and increasing DAUDA (0.02-12.7 µM). (**C and D**) The primary spectral components identified from SVD analysis for the reverse titration (C) and forward titration (D) are shown. In C, the primary component matched the emission peak of DAUDA-FABP1 (λ_em_ 509 nm, dark blue line). In D, the primary spectral component matched the emission peak of DAUDA in solution (λ_em_ 559 nm, dark blue line). Scree plots showing the relative strength (singular value) of the first 5 spectral components from SVD analysis of the reverse and forward titration spectra are shown in the insets. (**E and F**) Binding isotherms for the reverse (E) and forward (F) titrations. The dotted lines in F show the formation of singly and doubly bound DAUDA-FABP1 in the forward titration. Replicate titrations done on separate days are represented as different colored open circles. COPASI numerical fits of a single site binding model (E) and two-site sequential model (F) are shown as solid black lines.

The binding affinity of DAUDA to hFABP1 was first determined by ‘reverse’ titrations with a constant concentration (0.05 μM) of DAUDA and hFABP1 concentrations ranging from 0 to 1.7 µM (Figure 3A). Under these conditions, it is expected that the presence of doubly occupied hFABP1 is negligible. This is borne out by SVD analysis (Figure 3C), which shows primarily the monotonic increase of a species with emission maximum at 509 nm (corresponding to DAUDA bound to hFABP1) with only a small contribution from free DAUDA (λ_max_ = 559 nm). For this and subsequent spectral deconvolution, the basis spectrum for the DAUDA-FABP1 complex was determined from a sample of DAUDA with hFABP1 in excess (Supplemental Materials). The basis spectrum for free DAUDA in solution was also determined experimentally (Supplemental Figure S6B). Following spectral deconvolution, the quadratic binding equation was fit to the data of specific fluorescence of the DAUDA-FABP1 complex as a function of hFABP1 concentration yielding a DAUDA K_d_ of 0.20 (95% CI [0.15, 0.25]). These results were verified by fitting a numerical model of bimolecular association to the data which also yielded a K_d_ value of 0.2 µM. This K_d_ value corresponds to a single high affinity binding site of DAUDA with hFABP1.

The potential of multiple DAUDA molecules binding hFABP1 was then explored using ‘forward’ titrations with a constant concentration of hFABP1 (0.3 μM) and DAUDA concentrations ranging from 0 to 12.7 µM. Saturation was not achieved despite the highest DAUDA concentrations exceeding the K_d_ determined via the reverse titrations by over 50-fold (Figure 3). This suggests that multiple DAUDA molecules bind to hFABP1 simultaneously. However, only two spectral components were observed that made significant contributions to the observed signal in the SVD analysis (Figure 3), suggesting that the fluorescence spectrum of the doubly bound DAUDA-FABP1 complex is indistinguishable from a singly-bound complex. A sequential two-site binding numerical model was fit to the specific fluorescence of DAUDA-FABP1, with K_d,1_ fixed to the value obtained from reverse titrations. Since the specific fluorescence of DAUDA-FABP1 had not saturated even at 12.7 µM DAUDA, the fitting only allowed estimation of the lower bound of K_d,2_ (3.3 µM based on the 95% confidence interval).

To directly confirm the multiple DAUDA binding inferred from fluorescence titrations and determine the stoichiometry of DAUDA and hFABP1 complexes, native protein MS was used. Upon addition of one and two equivalents of DAUDA to hFABP1(10:10 and 20:10 µM), an *m/z* peak corresponding to apo-FABP1 was observed, as were additional higher-intensity peaks shifted to larger *m/z* values (Figure 4). Upon charge-state deconvolution, mass shifts of +434 and +868 Da corresponding to singly and doubly bound FABP1-DAUDA complexes, respectively, were observed supporting the findings from the fluorescence titrations. A considerable portion of apo-FABP1 was also observed which may be due to dissociation of the non-covalent DAUDA-FABP1 complexes in the electrospray process and in the gas phase.

**Figure 4:**
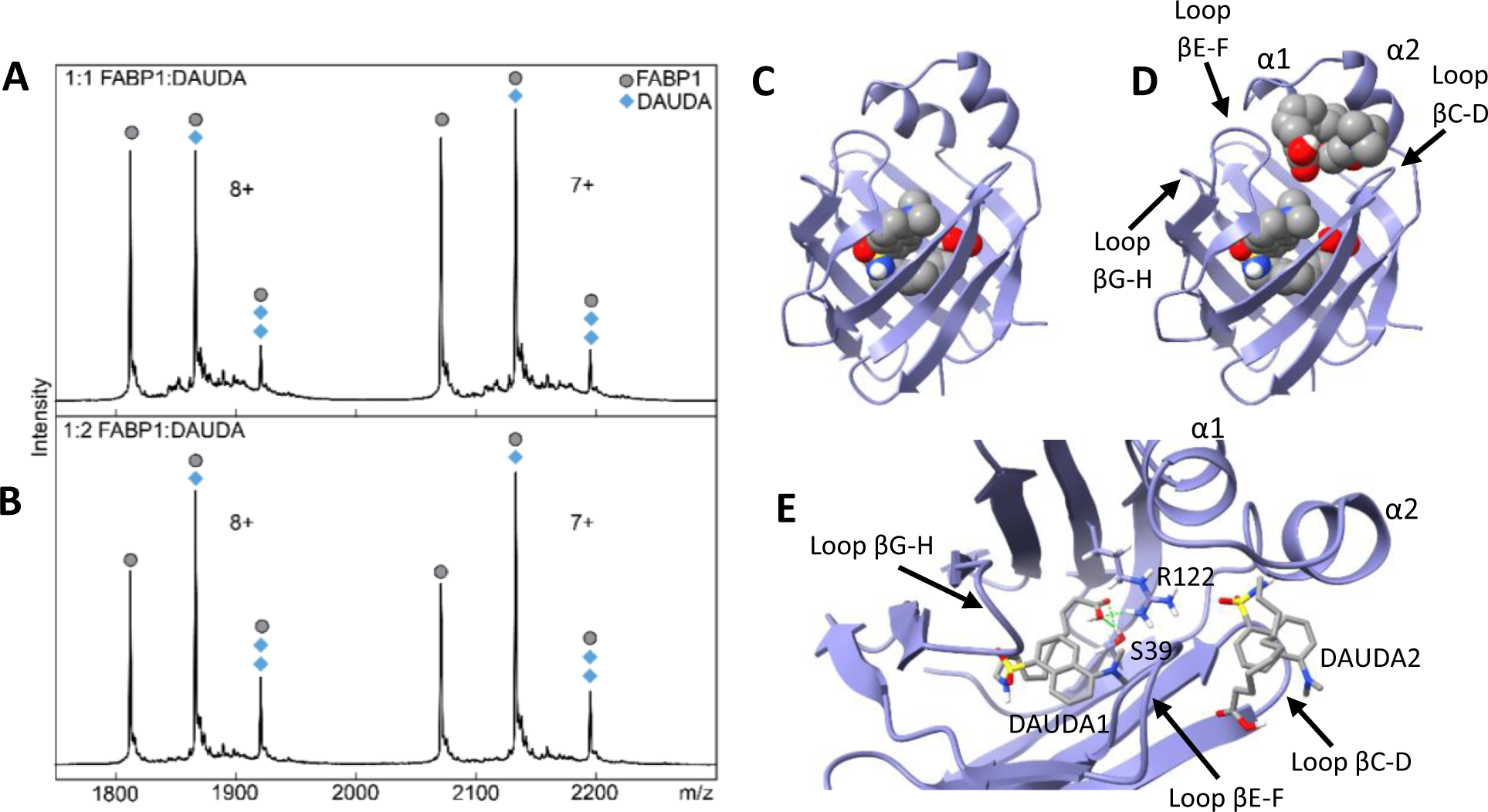
Native mass spectra and docking of DAUDA with hFABP1. Native mass spectra of a mixture of hFABP1 and DAUDA at **(A)** 1:1 (10:10 µM) and (**B)** 1:2 (10:20 µM) FABP1:DAUDA ratios. The mass of apo-FABP1 for each charge state (*n*+) is observed with additional *m/z-*shifted peaks corresponding to the molecular mass of 1 and 2 DAUDA (+434 and +868, respectively). Circle markers denote the apo form of the protein, while diamond markers denote peaks that correspond to FABP1-DAUDA and FABP1-2DAUDA complexes at the given charge state. (**C** and **D**) Structure of hFABP1 (PDB: 2LKK) docked with 1 (C) and 2 (D) molecules of DAUDA. (**E**) Top-down view of the docked structure in D. DAUDA1 corresponds to the DAUDA in center-bottom of the binding cavity. DAUDA1 interacts with sidechains from residues S39 and R122. DAUDA2 is located in the portal region in close proximity to the alpha helical domains.

To explore the binding modes of the two DAUDA within the binding cavity of hFABP1, two DAUDA were docked sequentially to hFABP1 (Figure 4C-E). The first DAUDA docked to the center-bottom of the hFABP1 binding cavity in a bent, U-shaped conformation with the DAUDA carboxyl group oriented toward R122 and S39 to form hydrogen bonds (ΔG_binding_ = - 8.05) (Figure 4C and 4E). This binding orientation was predicted to correspond to the high-affinity DAUDA binding site determined via fluorescence titrations and is consistent with the orientation of oleic acid (OA) (Cai *et al*., 2012) and palmitic acid (PA) (Sharma and Sharma, 2011) within hFABP1 determined from NMR and crystal structures. The second DAUDA was then docked sequentially and bound to a site near the portal region of hFABP1 (Figure 4D). Binding of the second DAUDA also resulted in a U-shaped conformation where both the carboxyl and dansyl groups of the molecule are oriented away from the hFABP1 binding cavity (ΔG_binding_ = -5.97) (Figure 4D and 4E). This binding site was predicted to correspond to the low affinity binding site of DAUDA detected in the fluorescence titrations.

### Arachidonic Acid as a Model Ligand for DAUDA Displacement Assays with hFABP1

AA is an endogenous fatty acid ligand for multiple FABPs including FABP1 (Veerkamp *et al*., 1999). AA was used as a model ligand to assess DAUDA displacement by ligands of hFABP1. The concentrations of DAUDA (0.5 µM) and hFABP1 (0.3 µM) used in these studies were chosen to keep DAUDA concentration low enough to ensure negligible formation of the doubly bound DAUDA-FABP1 complex. hFABP1 concentration was chosen to be as low as possible based on fluorescence assay sensitivity. Under these conditions, AA appeared to completely displace DAUDA fluorescence (Figure 5A). When hFABP1 binding was saturated with AA, the fluorescence spectrum resembled the spectrum of DAUDA free in solution. Similar to SVD analysis of reverse and forward titrations, only two spectral components were identified corresponding to DAUDA-FABP1 and DAUDA in solution (Figure 5B). Based on SVD analysis, the specific fluorescence of DAUDA in solution increased with the addition of AA (Figure 5C) consistent with DAUDA displacement from hFABP1 by AA. These results suggest a lack of a ternary DAUDA-AA-FABP1 complex formation, and that AA completely displaces DAUDA from hFABP1, with an apparent K_d_ of 0.08 ± 0.01 µM (Figure 5D, Table 1). These results are consistent with the K_i,app_ reported previously (0.11 µM) for AA in displacement assays using ANS (Huang *et al*., 2014).

**Figure 5:**
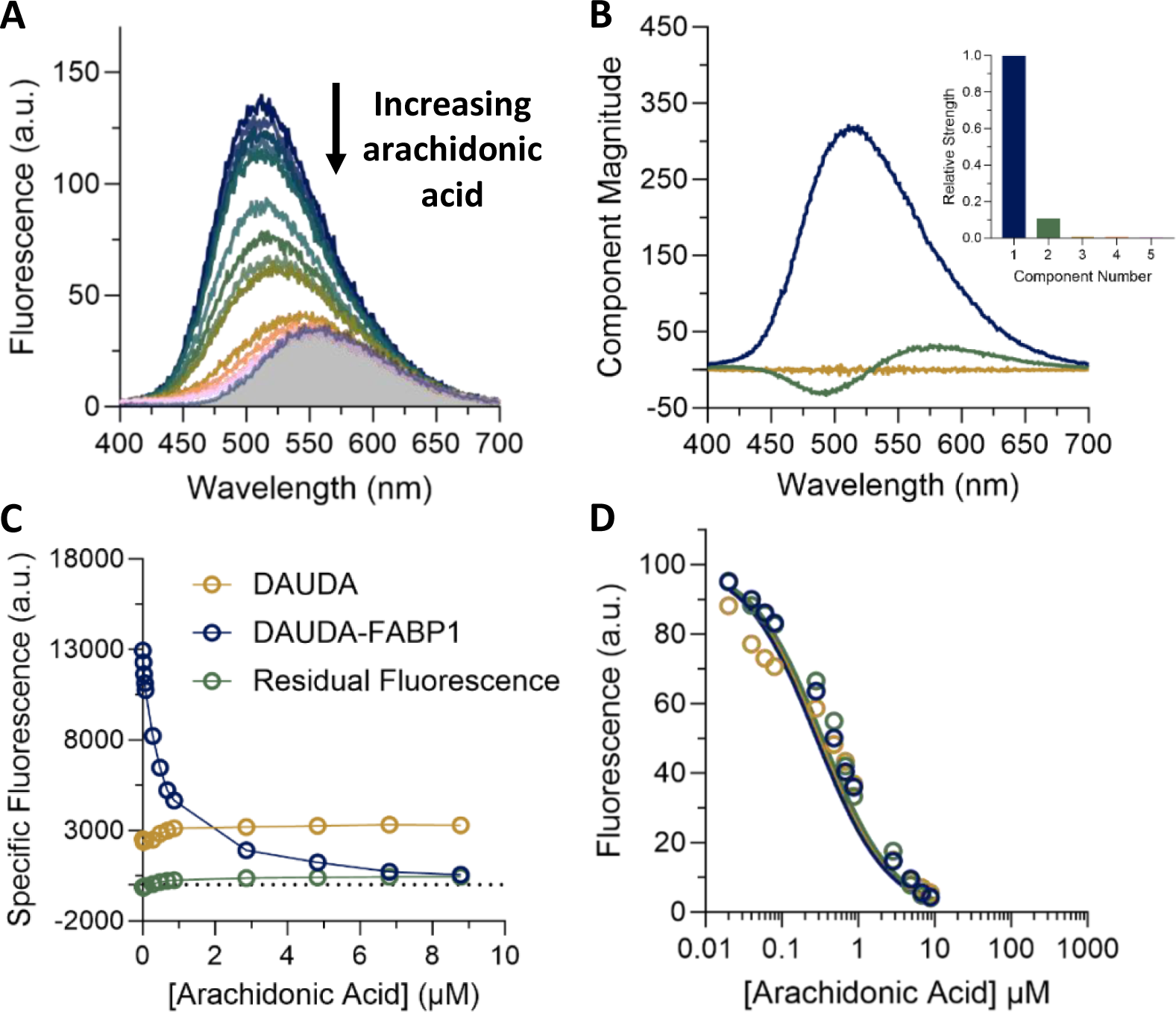
Displacement of DAUDA (0.5 µM) from FABP1 (0.3 µM) by increasing concentrations of arachidonic acid (0.02-8.8 µM). (**A**) Different colored spectra represent varying concentrations of arachidonic acid, increasing from top (dark blue, 0 µM) to bottom (pink, 8.8 µM). The shaded area is the spectrum of 0.5 µM unbound DAUDA in buffer. (**B**) The primary component (dark blue) identified from singular value decomposition (SVD) of the fluorescence spectra in A matches the emission peak for DAUDA-FABP1 (λ_em_ 509 nm). The second component (green) corresponds to changes in the fluorescence of DAUDA in solution. Scree plot showing the strength (singular value) of the first 5 spectral components from SVD analysis of the spectra in A is shown in the inset. (**C**) Relative change in the specific fluorescence of DAUDA in solution (gold), DAUDA bound to FABP1 (dark blue) with increasing arachidonic acid concentrations. (**D**) DAUDA displacement curve for arachidonic acid. A competitive binding model (solid lines) was fit to data for 3 replicate experiments done on separate days (dark blue, green and gold) to determine the K_d_ for arachidonic acid with hFABP1 (Table 1).

|  | **K<sub>d</sub>**<br>**(μM)** | **EC<sub>50</sub>**<br>**(μM)** | **F<sub>res</sub>**<br>**(% fluorescence remaining)** |
| --- | --- | --- | --- |
| Arachidonic acid | 0.08 ± 0.010 | 0.42 ± 0.11 | 2.2 ± 3.6 |
| Diazepam | N.D. | >50 | N.D. |
| Diclofenac | 2.5 ± 1.3 | 3.9 ± 2.1 | 15 ± 3.0 |
| R-flurbiprofen | N.D. | 4.4 ± 3.8 | 40 ± 1.6 |
| S-flurbiprofen | 2.2 ± 0.93 | 3.3 ± 1.4 | 48 ± 1.1 |
| Gemfibrozil | 3.6 ± 1.1 | 5.6 ± 1.7 | 18 ± 2.8 |
| Ibuprofen | 9.9 ± 0.60 | 15 ± 0.85 | 28 ± 4.9 |
| Pioglitazone | 1.0 ± 0.30 | 1.8 ± 0.47 | 5.2 ± 2.4 |
| Sulfaphenazole | 15 ± 1.7 | 21 ± 3.0 | N.D. |
| Tolbutamide | 20 ± 9.6 | 29 ± 14 | N.D. |

Values are reported as means ± S.D from 3 replicate experiments
N.D. – could not be determined with confidence

### Variety of Drug Ligands bind to hFABP1 and Form Ternary DAUDA-drug-FABP1 Complexes

The DAUDA displacement assay developed with AA was used to test the binding of diazepam, diclofenac, fluoxetine, flurbiprofen, gemfibrozil, ibuprofen, pioglitazone, sulfaphenazole and tolbutamide to hFABP1. All drugs tested except fluoxetine decreased DAUDA fluorescence, indicating these drugs bound to hFABP1 (Figure S7). Binding was confirmed via titrations using DAUDA displacement (Figure 6 and Figure S8). Kinetic modeling robustly yielded apparent K_d_ values for all drugs except (R)-flurbiprofen and diazepam. Therefore, the EC_50_ values are instead reported for these two ligands, with the caveat that this empirical parameter may conceal some more complex binding behavior. Diazepam, in particular did not achieve saturation in the tested concentration range, and so the uncertainty in EC_50_ is high. The apparent affinity values for most of the drugs characterized were within low micromolar range (Table 1), consistent with the high binding promiscuity of hFABP1.

**Figure 6:**
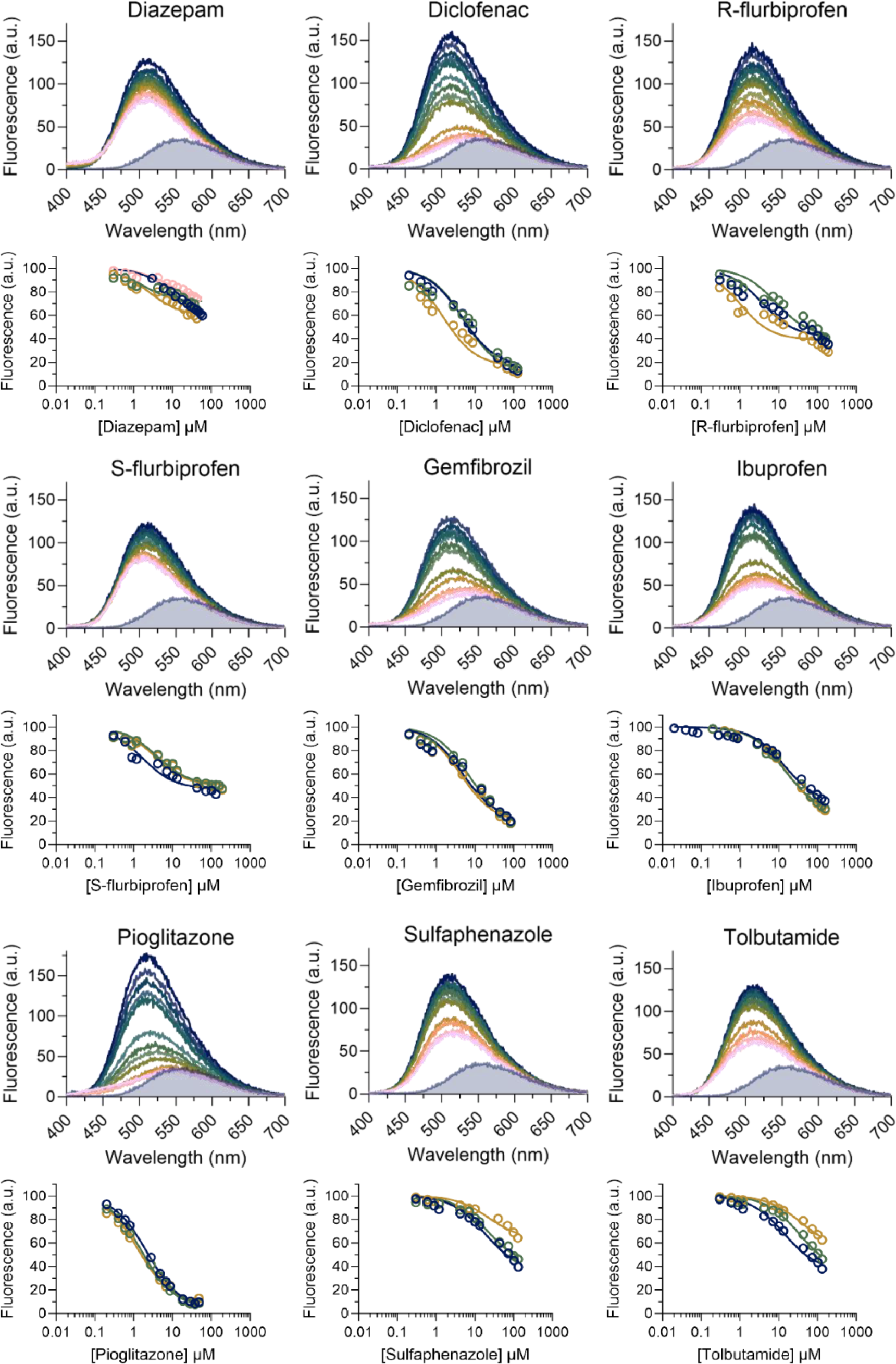
DAUDA displacement from FABP1 by drug ligands. The fluorescence spectra for DAUDA displacement titrations with diazepam, diclofenac, R- and S-flurbiprofen, gemfibrozil, ibuprofen, pioglitazone, sulfaphenazole and tolbutamide as ligands are shown across the rows. In each titration spectra, the top dark blue spectrum represents DAUDA (0.5 µM) prebound with FABP1 (0.3 µM) in the absence of ligand and each subsequent colored spectrum represents increasing concentrations of ligand. The shaded area is the spectrum of 0.5 µM unbound DAUDA in the absence of FABP1 and ligand. Corresponding DAUDA displacement curves are shown below each spectra. A ternary complex binding model was fit to replicate experiments done on separate days (dark blue, green, gold and pink circles) to determine K_d_ values summarized in Table 1.

None of the drugs completely eliminated the fluorescence of DAUDA to the levels observed for free DAUDA in solution (gray shaded spectrum in Figure 6). Diclofenac, gemfibrozil and pioglitazone decreased the fluorescence of DAUDA-FABP1 by >82 % at saturation (i.e., *F_res_* <18%), while the maximal decrease for (R)- and (S)-flurbiprofen and ibuprofen ranged between 52-72% (Table 1). The *F_res_* values for diazepam, sulfaphenazole and tolbutamide could not be determined with confidence. Inspection of the fluorescence spectra with drug ligands showed that for all tested drugs the spectra at saturation were blue shifted relative to the spectrum of free DAUDA in solution (Figure 6). For all ligands tested, the primary spectral component matched the expected λ_em_ of DAUDA-FABP1 (509 nm) (Figure S8). For the majority of ligands, the second spectral component corresponded to the fluorescence of DAUDA in solution. However, unlike titrations with AA, the specific fluorescence of DAUDA in solution did not increase with increasing concentrations of drug (Figure S8). Taken together these findings suggest that, unlike AA which appeared to completely displace DAUDA from hFABP1, drug ligands bound to hFABP1 simultaneously with DAUDA as a ternary complex altering the fluorescence characteristics of DAUDA-FABP1. In support of the presence of such ternary complexes, SVD analysis identified unique spectral components that were different than DAUDA alone in solution or DAUDA-FABP1 in (R)- and (S)-flurbiprofen titrations (Figure S10). The SVD analysis could not, however, consistently identify the presence of such species that were spectrally distinct in other titrations.

Native protein mass spectrometry was used to directly detect ternary DAUDA-diclofenac-FABP1 complexes (Figure 7). The native MS showed *m/z* shifts corresponding to DAUDA-FABP1, diclofenac-FABP1, ternary DAUDA-diclofenac-FABP1 and ternary DAUDA-FABP1-DAUDA complexes. High intensity peaks were also observed for apo-FABP1. No peaks were observed corresponding to hFABP1 bound with 2 diclofenac molecules under these experimental conditions. Consistent with the higher binding affinity of DAUDA in comparison to diclofenac with hFABP1, the majority of hFABP1 was in complex with DAUDA with a small fraction of hFABP1 found complexed with diclofenac.

**Figure 7:**
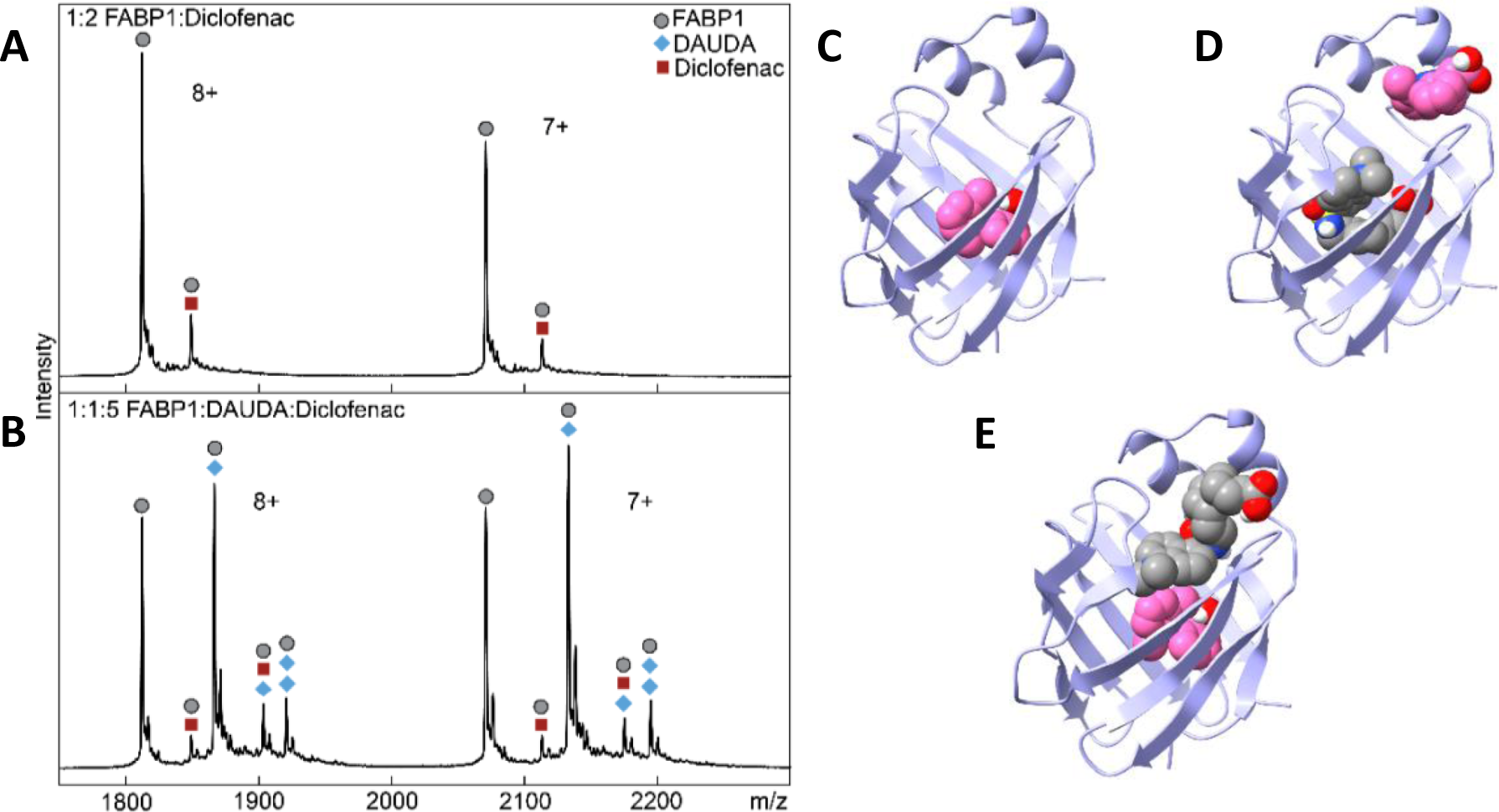
Characterization of diclofenac binding to hFABP1 via native protein mass spectrometry and molecular docking. **(A**) Native mass spectrum of hFABP1 and diclofenac at a 1:2 hFABP1:diclofenac ratio. Circle markers represent the apo form of the protein, while square markers denote *m/z*-shifted peaks that correspond with diclofenac. **(B**) Native mass spectrum of hFABP1, DAUDA, and diclofenac at 1:1:5 ratios, which uses the same marker labels as A, with the addition of diamond markers that denote *m/z*-shifted peaks corresponding to the association of DAUDA. (**C-E**) Docked structures of hFABP1 complexes showing potential binding orientations of singly bound diclofenac (C, pink) and diclofenac in complex with DAUDA (D and E, gray).

The potential binding orientation of diclofenac when in complex with DAUDA and hFABP1 were assessed via molecular docking (Figure 7C-E). Similar to DAUDA, the carboxyl head group of singly bound diclofenac was oriented toward S39 and R122 centered within the hFABP1 binding cavity and interacted via hydrogen bonding (ΔG_binding_ = -7.2) (Figure 7C). Since diclofenac decreased DAUDA fluorescence by ∼90% but the free DAUDA signal did not increase proportionately and the resulting blue shifted spectrum indicated a ternary DAUDA-FABP1-diclofenac complex, we explored the possibility of sequential binding modes of DAUDA and diclofenac. Sequential docking studies were performed where either DAUDA or diclofenac were first docked to hFABP1 before subsequent docking of the other ligand (Figure 7D and 7E). With DAUDA in the hFABP1 binding cavity, diclofenac bound to a site near the portal region where the carboxyl group was oriented away from the binding cavity interacting with residues K31 and S56 (ΔG_binding_ = -6.5). When DAUDA was docked with diclofenac in the binding cavity, DAUDA adopted an elongated “head out” conformation (Figure 7E). The carboxyl head group of DAUDA was oriented near the portal domain facing away from the binding cavity while the dansyl group was buried within the hFABP1 binding cavity (ΔG_binding_ = -7.2). This was in contrast to the U-shape confirmation of the second DAUDA resulting from sequential docking of two DAUDA (Figure 4D).

Distinct spectral components in (R)- and (S)-flurbiprofen titration spectra identified by SVD analysis indicated the formation of ternary flurbiprofen-DAUDA-FABP1 complexes. Hence, docking studies were also performed with (R)- and (S)-flurbiprofen to explore the potential binding orientations of flurbiprofen in complex with DAUDA and FABP1. Both singly docked (R)- and (S)-flurbiprofen (Figure 8A and 8B, respectively) had similar orientations within the FABP1 binding cavity (ΔG_binding_ = -7.6 and -7.5, respectively). The carboxyl groups for both molecules interacted with residues S39, S134 and R122 via hydrogen bonding. When (R)- and (S)-flurbiprofen were independently docked to hFABP1 with DAUDA present (Figure 8C and 8D), both flurbiprofen molecules were positioned near the portal region and α-helical domain of hFABP1. However, (R)-flurbiprofen bound further in the hFABP1 binding cavity than (S)-flurbiprofen and was closer in proximity to DAUDA (ΔG_binding_ = -6.9) (Figure 8E). In contrast, (S)-flurbiprofen bound within the opening of the portal region where the carboxyl group interacted with residues K31 and S56 (ΔG_binding_ = -6.8).

**Figure 8:**
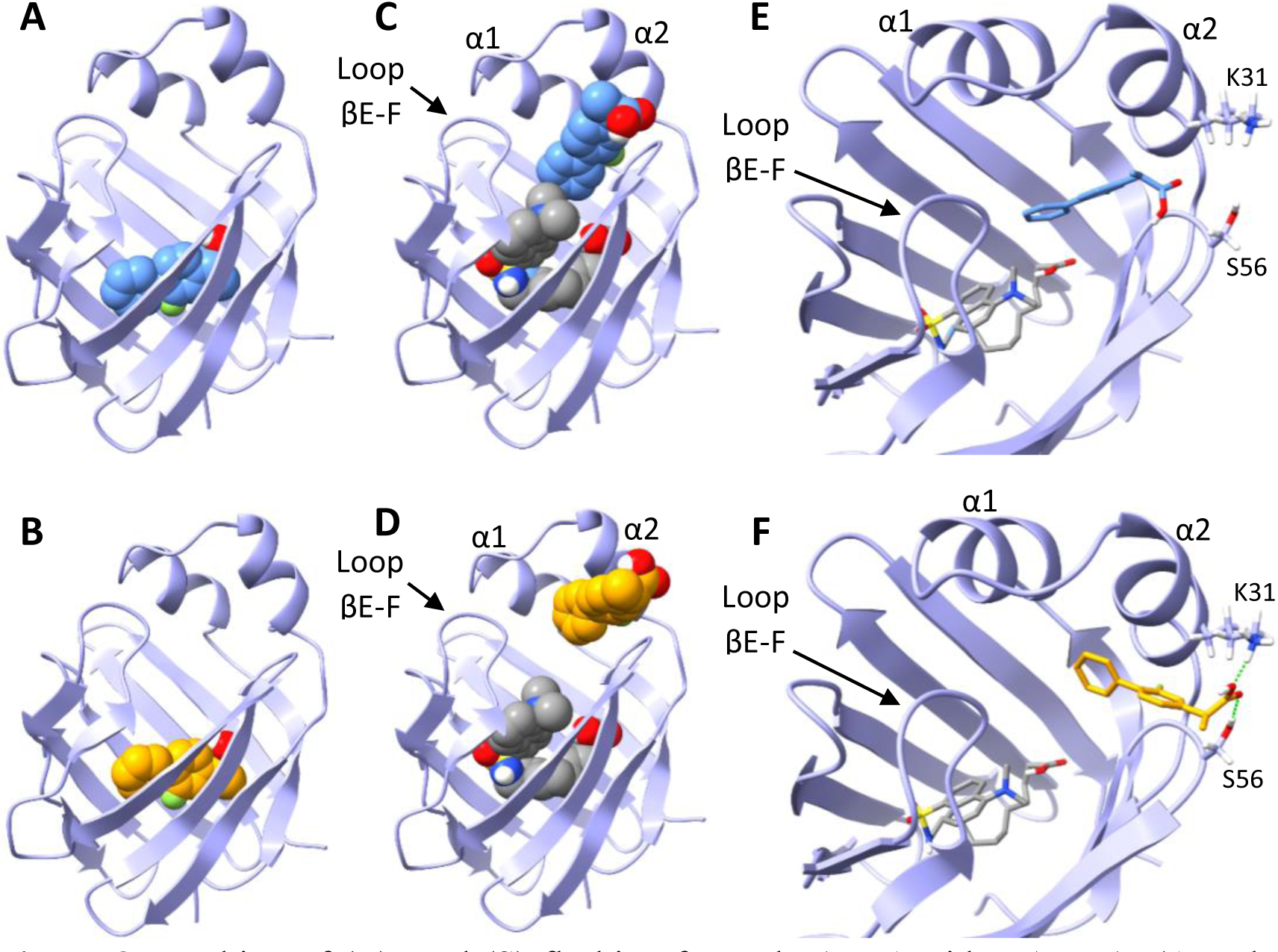
Docking of (R)- and (S)-flurbiprofen to hFABP1 with DAUDA. (**A** and **B**) Potential binding orientations of singly bound (R)-flurbiprofen (A, blue) and (S)-flurbiprofen (B, orange) in the absence of DAUDA. (**C** and **D**) Binding orientations of (R)- and (S)-flurbiprofen are shown in the presence of DAUDA (gray) in the hFABP1 binding cavity. **(E** and **F**) Molecular stick models showing the distinct positions of (R)-(E) and (S)-flurbiprofen (F) at the portal domain of hFABP1 in the presence of DAUDA in the binding cavity. Predicted hydrogen bonding for the carboxyl group of (S)-flurbiprofen is shown in green.

### hFABP1 Binding Alters 4’-OH-diclofenac Formation Kinetics by CYP2C9

To determine the effect of hFABP1 on diclofenac metabolism by CYP2C9, the formation kinetics of 4’-OH-diclofenac by recombinant CYP2C9 was characterized in the presence and absence of 20 µM hFABP1 (Figure S11). At this hFABP1 concentration, a significant portion of diclofenac is expected to be bound to hFABP1 based on the Kd value determined for diclofenac using DAUDA displacement assay. The apparent nominal Km for 4’-OH-diclofenac formation by CYP2C9 was significantly higher in the presence of hFABP1 (5.8 ± 1.5 µM) compared to in the absence of hFABP1 (1.4 ± 0.2 µM) (Table 2, Figure S11). This suggests that hFABP1 sequesters diclofenac from CYP2C9 mediated metabolism. Surprisingly, the apparent k_cat_ was decreased in the presence of hFABP1 when compared to the incubations done in the absence of hFABP1 (Figure S11).

**Table 2:**
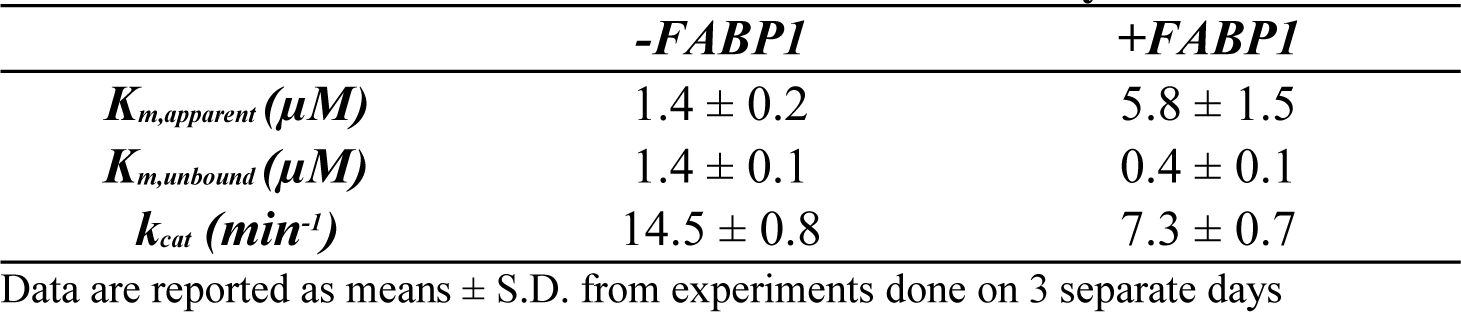
Kinetics of 4’-OH-diclofenac Formation by CYP2C9.

To test whether the effect of hFABP1 on diclofenac Km could be explained by the free drug hypothesis, unbound concentrations of diclofenac were determined in the incubations with and without hFABP1. The mean diclofenac fu in the absence of FABP1 was 1.0 ± 0.04 while the fu in the presence of 20 μM FABP1 ranged from 0.1-0.5 and was diclofenac concentration dependent (Figure 9). Based on the free concentrations of diclofenac determined in these experiments, the K_m,u_ and the k_cat_ in the presence of hFABP1 for 4’-OH-diclofenac formation were 0.4 ± 0.1 μM and 7.3 ± 0.7 min^-1^, respectively (Figure 9, Table 2). The k_cat_ value was significantly lower (*p* < 0.0005) in the presence of hFABP1 (7.3 ± 0.7 min^-1^) than in the absence of hFABP1 (14.5 ± 0.8 min^-1^). A trend towards a decrease in K_m,u_ was observed but not significantly different (0.4 ± 0.1 and 1.4 ± 0.1 μM, in the presence and absence of hFABP1, respectively). These data suggest that in addition to sequestering and binding diclofenac, hFABP1 directly interacts with CYP2C9 to noncompetitively inhibit diclofenac metabolism and CYP2C9 catalytic activity.

**Figure 9:**
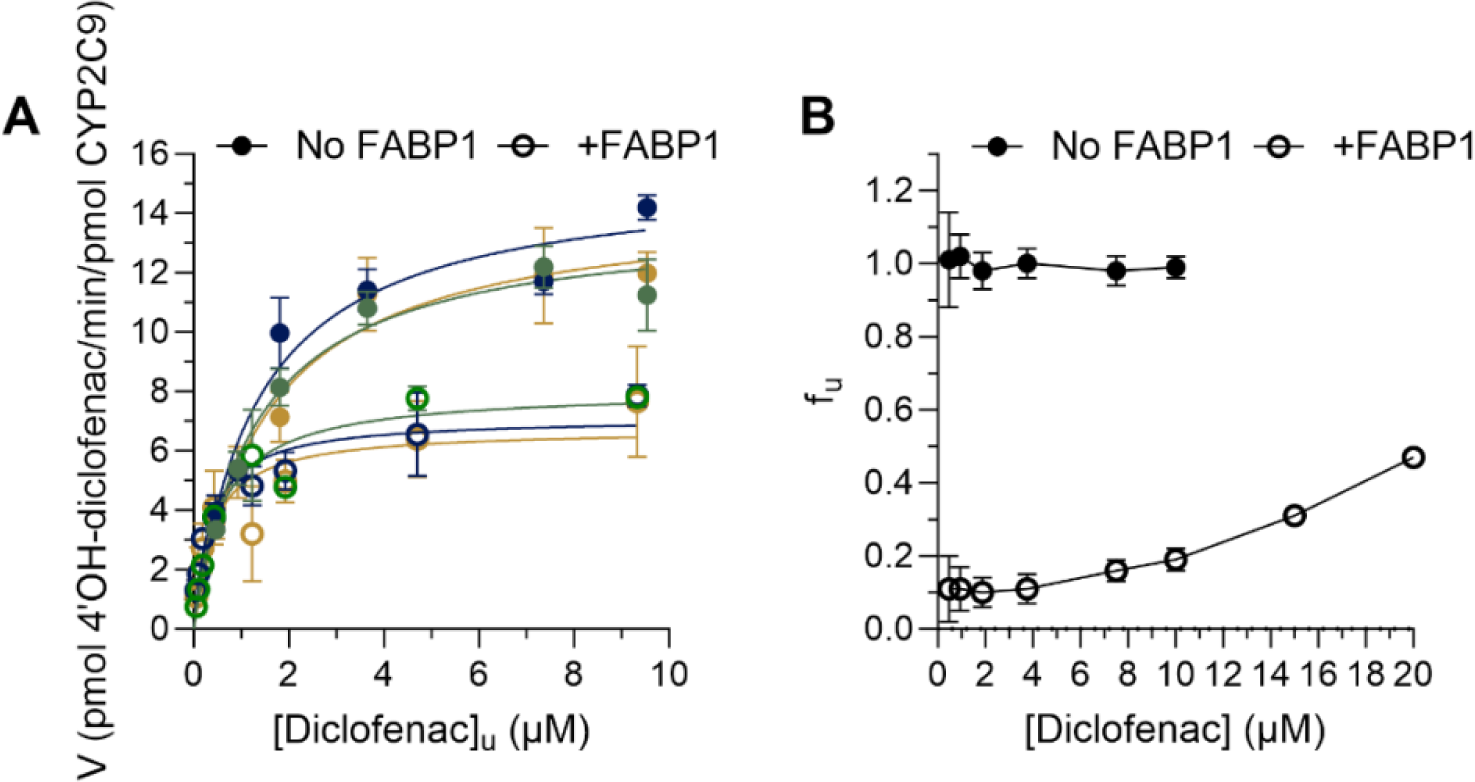
Impact of FABP1 on 4’-OH-diclofenac formation kinetics by CYP2C9. **(A)** 4’-OH-diclofenac formation by CYP2C9 in the presence (open circles) and absence (solid circles) of 20 µM FABP1 is shown. Paired replicate experiments from three separate days are shown in dark blue, green and gold**. (B)** Free fraction (fu) of diclofenac for all nominal concentrations used in kinetic experiments with FABP1 (20 μM) was determined using magnetic silica beads and magnetic Ni-NTA agarose beads as described in Materials and Methods. The unbound Km and kcat values for diclofenac are summarized in Table 2.

## Discussion

hFABP1 is highly abundant in the liver and intestines and serves as a major binding protein for lipophilic compounds. Yet, drug binding to hFABP1 and the role of hFABP1 in drug distribution and metabolism have been poorly defined. *In vivo*, hFABP1 is likely present as a mixture of apo-FABP1 and endogenous lipid bound holo-FABP1 (Schroeder *et al*., 1998). The binding capacity of FABP1 *in vivo* is increased by the possibility of two ligands binding simultaneously to FABP1 as shown for oleate and palmitate using native MS (Santambrogio *et al*., 2013), NMR (Cai *et al*., 2012) and crystallography (Sharma and Sharma, 2011). Drug binding to FABP1 can be complicated as drugs may bind to apo-FABP1 or as a second ligand to FABP1 already bound with fatty acids. To probe these drug binding modalities, DAUDA was chosen as the fluorescent ligand in this study. DAUDA is a well characterized fatty acid derivative that is a larger ligand than the commonly used ANS and binds FABP1 with higher affinity (Thumser and Wilton, 1994; Davies *et al*., 2002; Luebker *et al*., 2002a; Norris and Spector, 2002). Hence, DAUDA binding likely mimics native lipid binding to FABP1.

Fluorescence titrations and native MS studies, together with the docking studies, support the conclusion that DAUDA binding captures the two binding sites of endogenous ligands with hFABP1. Based on fluorescence titrations, DAUDA has a high (K_d1_=0.2µM) and a low (K_d2_>3.3µM) affinity binding site on hFABP1 that likely correspond to the binding sites of endogenous fatty acids. Indeed, the K_d_ values for DAUDA are comparable to the high (0.009-0.2µM) and low (0.06-7.6µM) affinity binding sites determined for OA with bovine and rFABP1 and PA for rFABP1 (Richieri *et al*., 1994, 1996; Rolf *et al*., 1995; Santambrogio *et al*., 2013). In docking studies, DAUDA bound within the bottom-center of the hFABP1 binding cavity in a U-shape orientation consistent with the orientation of OA and PA in the high affinity binding site (Sharma and Sharma, 2011; Cai *et al*., 2012). Docking of a second DAUDA resulted in DAUDA binding near the portal region where a second putative low affinity binding was also reported.

The use of DAUDA in fluorescence displacement assays can be challenging due to the background fluorescence of DAUDA which interferes with direct measurements of DAUDA-FABP1 fluorescence. Background correction methods subtracting free DAUDA fluorescence have been reported (Thumser *et al*., 1996; Davies *et al*., 2002; Luebker *et al*., 2002b; Elmes *et al*., 2019). These methods do not account for the different free DAUDA concentrations in the presence and absence of FABP1 or that multiple DAUDA may bind to FABP1 simultaneously (Norris and Spector, 2002). To address these concerns, SVD analysis was introduced here to determine the specific contribution of DAUDA-FABP1 to observed fluorescence spectra. The SVD-based spectral deconvolution allowed separation of the DAUDA-FABP1 signal from fluorescence due to free DAUDA, enabling rigorous ligand binding analysis using the DAUDA displacement assay. Of the drugs studied here and found to bind to hFABP1, diazepam, diclofenac, flurbiprofen, gemfibrozil and ibuprofen have been previously shown to bind to rFABP1 (Chuang *et al*., 2008). Their binding to rFABP1 was measured based on ANS fluorescence displacement, and a two-site competition model was fit to the data. For all five drugs two binding sites in rFABP1 were kinetically detected with high affinity binding K_i_s ranging from 1 to 47µM and low affinity binding K_i_s being 10-200-fold higher, 35-448µM (Chuang *et al*., 2008). The K_d_ values determined here for diclofenac, flurbiprofen, gemfibrozil and ibuprofen with hFABP1 using DAUDA displacement and SVD analysis ranged from 2 to 10µM. The apparent EC_50_ for diazepam with hFABP1 was >50µM. K_d_ values for diclofenac, (S)-flurbiprofen and gemfibrozil for hFABP1 in this study were within 2-fold of the high affinity K_i_ values reported for rFABP1. However, the affinity of ibuprofen was ∼5 times greater for hFABP1 than rFABP1, and the affinity of diazepam was at least 2 orders of magnitude lower for hFABP1 than rFABP1. Although the potential for multiple binding sites and formation of ternary complexes complicates the comparison and interpretation of these affinity values, these data suggest that while drug binding characteristics for rFABP1 and hFABP1 are qualitatively similar, drug binding data with rFABP1 does not translate quantitatively to hFABP1.

Previous NMR studies identified two different binding sites in rFABP1 for ANS, ketorolac and ibuprofen (Chuang *et al*., 2008). Residues located in the bottom of the FABP1 β-barrel were perturbed by ligand binding. In the presence of ligand concentrations in 2-fold excess of the protein concentration, additional residues were perturbed in the portal region. These findings are consistent with the high affinity binding site for drugs in the bottom of the β-barrel and the low affinity binding site in the portal region. The data collected in this study with fluorescence displacement of DAUDA, native MS and docking studies with hFABP1 support similar binding characteristics with hFABP1 with drug molecules occupying one of the two binding sites with DAUDA occupying the other simultaneously.

Fluorescence data suggest that all drugs tested here form ternary complexes with DAUDA and hFABP1. *F_res_* values at saturation ranged from 5 to 48% between drugs suggesting that in contrast to AA, drug ligands do not completely displace DAUDA from hFABP1 but rather bind to hFABP1 simultaneously with DAUDA. The formation of ternary complexes was supported by the observation of a clear blue shift in the fluorescence spectrum of DAUDA in the presence of drug ligands and by the lack of increase in the fluorescence signal of DAUDA free in solution in the titrations (Figure 6). Ternary complex formation was confirmed with diclofenac by native MS where both diclofenac and DAUDA bound to hFABP1 simultaneously.

One may speculate that drugs (diclofenac, pioglitazone, gemfibrozil) that decrease DAUDA fluorescence almost completely may bind in the high affinity site in hFABP1 while the drugs ((R)- and (S)-flurbiprofen and ibuprofen) that decrease DAUDA fluorescence by only 52-72% bind in the α-helical lid region resulting in different fluorescence spectra at saturation due to the different orientation of DAUDA within hFABP1. Indeed, SVD analysis did not identify unique spectral components with diclofenac but did so with (R)- and (S)-flurbiprofen. Docking studies where DAUDA was sequentially docked to hFABP1 with diclofenac in the binding cavity showed DAUDA binding in a head out position near the portal region of hFABP1, supporting the hypothesis that DAUDA binds to the low affinity site in the presence of diclofenac. Sequential docking studies with DAUDA bound to hFABP1 placed (R)- and (S)-flurbiprofen at the portal region with distinct binding orientations possibly explaining differences in the fluorescence spectra and unique spectral components identified for the enantiomers. A limitation of the analysis presented here is that it does not account for potential changes in DAUDA binding affinity due to the presence of other hFABP ligands. Further studies are needed to explore the drug binding with hFABP1 and how different binding orientations and conformations alter drug metabolism and disposition as well as lipid metabolism and signaling.

The impact of drug binding to apo-hFABP1 on drug metabolism by CYPs was evaluated using diclofenac metabolism by CYP2C9 as a model reaction. The K_m,u_ and k_cat_ of 4’OH-diclofenac formation were decreased in the presence of hFABP1 by ≥50%, suggesting that hFABP1 interacts with CYP2C9 via a protein-protein interaction and modulates metabolism by CYP2C9. FABP1 has been proposed to deliver ligands directly to lipid membranes (Davies *et al*., 2002), but another report suggests FABP1 releases its ligand into solution rather than interacting with lipid membranes (Hsu and Storch, 1996). FABP1 has also been found to directly interact with metabolic enzymes and receptors to modulate their activity. FABP1 interactions with CPTI enhanced activity toward LCFA-CoA (Hostetler *et al*., 2011), and interactions with PPARα were proposed as a mechanism for PPARα activation (Hostetler *et al*., 2009). Similar protein-protein interactions likely occur with CYP2C9. The observations of the effects of hFABP1 on CYP2C9 activity are similar to those observed previously with other intracellular lipid binding protein family members, cellular retinoic acid binding proteins, and CYP26 enzymes (Nelson *et al*., 2016; Zhong *et al*., 2018; Yabut and Isoherranen, 2022). This suggests that binding proteins may be more promiscuous modulators of cellular metabolism than previously described.

The results shown here unequivocally establish that many drugs bind to hFABP1 and strongly suggest that hFABP1 binding will alter drug distribution and metabolism in the human liver. These findings have important implications for modeling drug disposition in the liver and for predicting drug clearance for drugs that bind to hFABP1. The formation of drug-DAUDA-FABP1 complexes suggests that drug ligands may not have to compete with endogenous ligands for hFABP1 binding but rather that in the human liver drugs may bind to hFABP1 as a ternary complex with an endogenous lipid. However, this is likely drug dependent as the binding modes of different drugs to hFABP1 bound with DAUDA likely vary as suggested by fluorescence and docking results for diclofenac and flurbiprofen. Future studies are needed with mixed drug-lipid-FABP1 complexes to fully unravel the role of hFABP1 in modulating drug metabolism.

## Supporting information

Biorender license for Fig2A

Biorender license For Fig S9

## Abbreviations

AA: Arachidonic acid
ANS: 8-anilinonaphthalene-1-sulfonic acid
CYP: cytochrome P450
DAUDA: 11-(Dansylamino)undecanoic Acid
FABP: fatty acid binding protein
*f_u_*: unbound fraction
LCFA: long chain fatty acid
LC-MS/MS: Liquid chromatography-tandem mass spectrometry
MGSB: magnetic silica beads
MS: Mass spectrometry
K_d_: equilibrium binding affinity constant
OA: oleic acid; PA: palmitic acid
SVD: singular value decomposition

## Acknowledgments

Figure 2A and Figure S9 were created with BioRender.com shapes.

## Data Availability Statement

All the research data supporting the results are reported in the main manuscript or supplemental materials. Raw data is available from the corresponding author upon request.

## Authorship Contributions

*Participated in research design:* Yabut, Martynova, Nath, Zercher, Bush, Isoherranen

*Conducted experiments:* Yabut, Martynova, Zercher

*Performed data analysis:* Yabut, Martynova, Nath, Zercher

*Wrote or contributed to the writing of the manuscript:* Yabut, Martynova, Nath, Zercher, Bush, Isoherranen

## Footnotes

No author has an actual or perceived conflict of interest with the contents of this article. This work was supported in part by grants from the National Institutes of Health [Grant T32 GM007750] and [Grant P01 DA032507]. NI is supported in part by the Milo Gibaldi Endowed Chair of Pharmaceutics to Department of Pharmaceutics, University of Washington

