## Supplemental materials for "Drugs Form Ternary Complexes with Human Liver Fatty Acid Binding Protein (FABP1) and FABP1 Binding Alters Drug Metabolism"

### Cloning and Expression of FABP1

The human FABP1 open reading frame was purchased from Origene (Rockville, MD, Cat. No. SC119222) and cloned into pET28a+ with an N-terminal hexa-histidine (6xHis) tag and a thrombin cleavage site using N-terminal NdeI and C-terminal HindIII restriction sites. The plasmid map and coding sequence are shown in Figure S1.

The expression construct was transformed into Rosetta 2 *E. coli* (Novagen, Madison, WI) and a freshly transformed colony was inoculated into a 25 mL starter culture of Luria Broth (LB) with kanamycin (50 µg/mL) and grown for 6 hours at 37°C in a shaking incubator at 250 rpm. 15 mL of starter culture was used to inoculate 1 L of LB-kanamycin which was then grown to an OD600 ≈ 0.6. The culture was cooled at room temperature for 20 minutes prior to adding 0.1 mM IPTG to induce FABP1 expression (Figure S2A). The protein was expressed at 18°C for 18 hours in a shaking incubator at 250 rpm. Cells were harvested by centrifugation at 4,000 g for 20 minutes then washed with phosphate buffered saline (PBS) containing 1 mM PMSF, pelleted, decanted and stored at -80°C until purification.

Thrombin Cleavage of the His tag: The thrombin cleavage protocol was optimized by testing cleavage at different combinations of thrombin concentrations (0.02-0.1 U per 1 µg his-tagged FABP1), incubations times (0.5-6 hours) and incubation temperatures (4, 21 and 37°C) in elution buffer (20 mM Tris pH 7.4 at 4°C with 500 mM NaCl and 400 mM imidazole) or elution buffer diluted with an equivalent volume of 20 mM Tris pH 7.4 at 4°C with 100 mM NaCl. Cleavage of the his-tagged FABP1 was assessed by SDS-PAGE and Coomassie staining. To streamline the purification protocol, an incubation time of 1 hour at 37°C was chosen which resulted in complete cleavage of his-tagged FABP1 using 0.03 U thrombin per 1 µg FABP1 in diluted elution buffer (20 mM Tris pH 7.4 at 4°C with 300 mM NaCl and 200 mM imidazole) (Figure S2C).

### Optimization of Purification and Delipidation of FABP1

Nickel purification of 6xHis FABP1: Frozen pellets were thawed on ice in lysis buffer (20 mM Tris pH 7.4 at 4°C, 500 mM NaCl, 30 mM imidazole) with protease inhibitor cocktail (Roche, cOmplete Mini EDTA-free), 1 mM PMSF and 25 U of benzonase. 1 mg/mL lysozyme was added and the cells were rocked on ice for 30 minutes. Cells were sonicated at 75% power for 30 seconds with a 1-minute rest on ice for 5 rounds (Figure S2B). The lysate was cleared at 20,000 g for 30 minutes and the supernatant filtered through a 0.22 µm syringe filter (Minisart, Sartorius, Göttingen, Germany) (Figure S2B). The filtered lysate was loaded onto a 90 mL Dynaloop (Bio-Rad, Hercules, CA) coupled to a DuoFlow fast protein liquid chromatograph (FPLC) (Bio-Rad, Hercules, CA) and run over a 1 mL HisTrap HP affinity column (GE Healthcare, Chicago, IL) equilibrated in lysis buffer at a flow rate of 0.75 mL/min. The column was washed with 10 column volumes of wash buffer (20 mM Tris pH 7.4 at 4°C, 500 mM NaCl, 30 mM imidazole) (Figure S2B) and FABP1 eluted using elution buffer (20 mM Tris pH 7.4 at 4°C, 500 mM NaCl) with a step gradient of increasing concentrations of imidazole (0-500 mM) in 1 mL fractions over 10 column volumes. The eluted protein was detected at 280 nm absorbance and the peak FABP1 containing fractions were pooled for subsequent steps of purification.

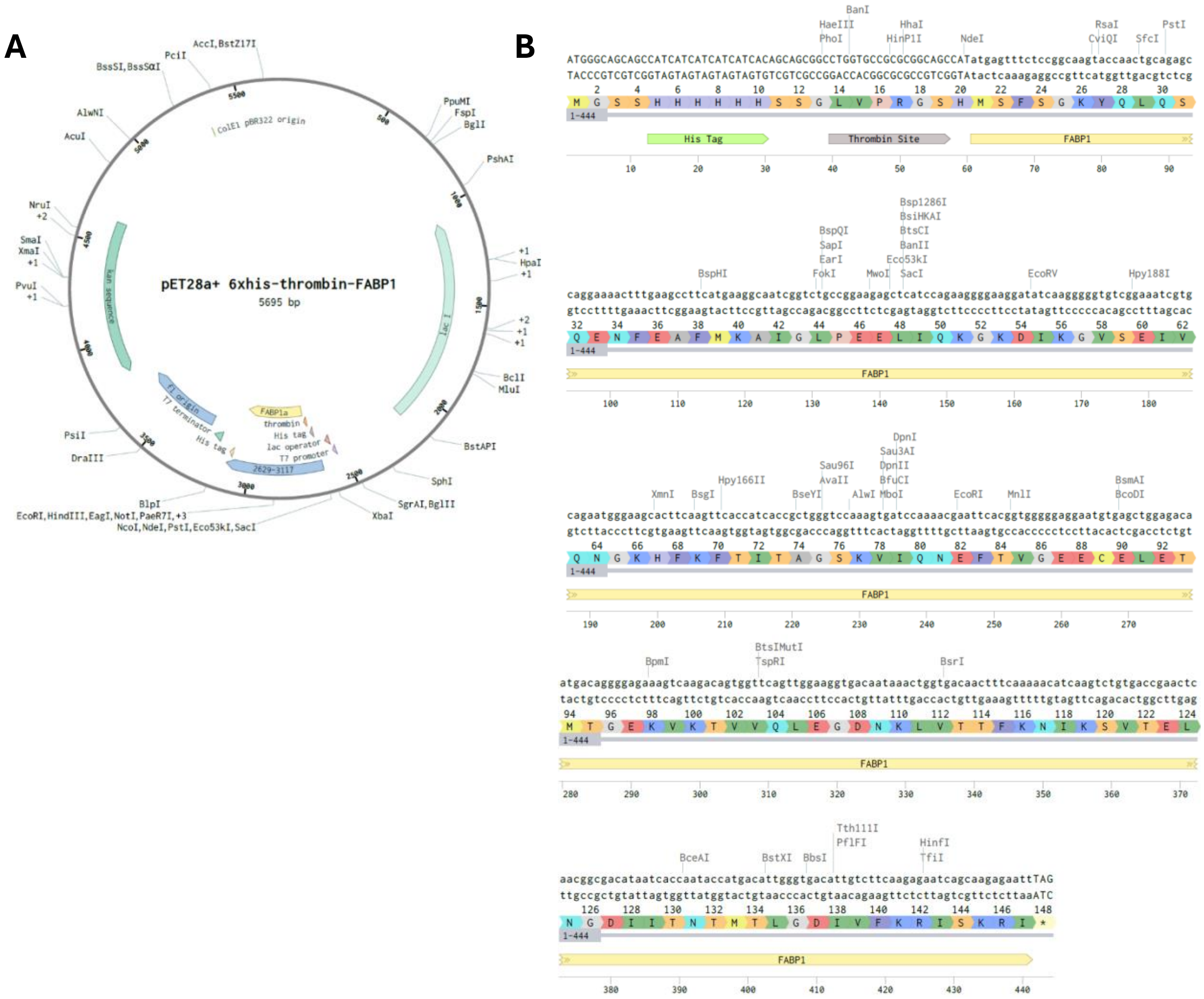


**Figure S1:** Plasmid map and coding sequence for recombinant FABP1 visualized by Benchling [Biology Software]. (2023). Retrieved from https://benchling.com.

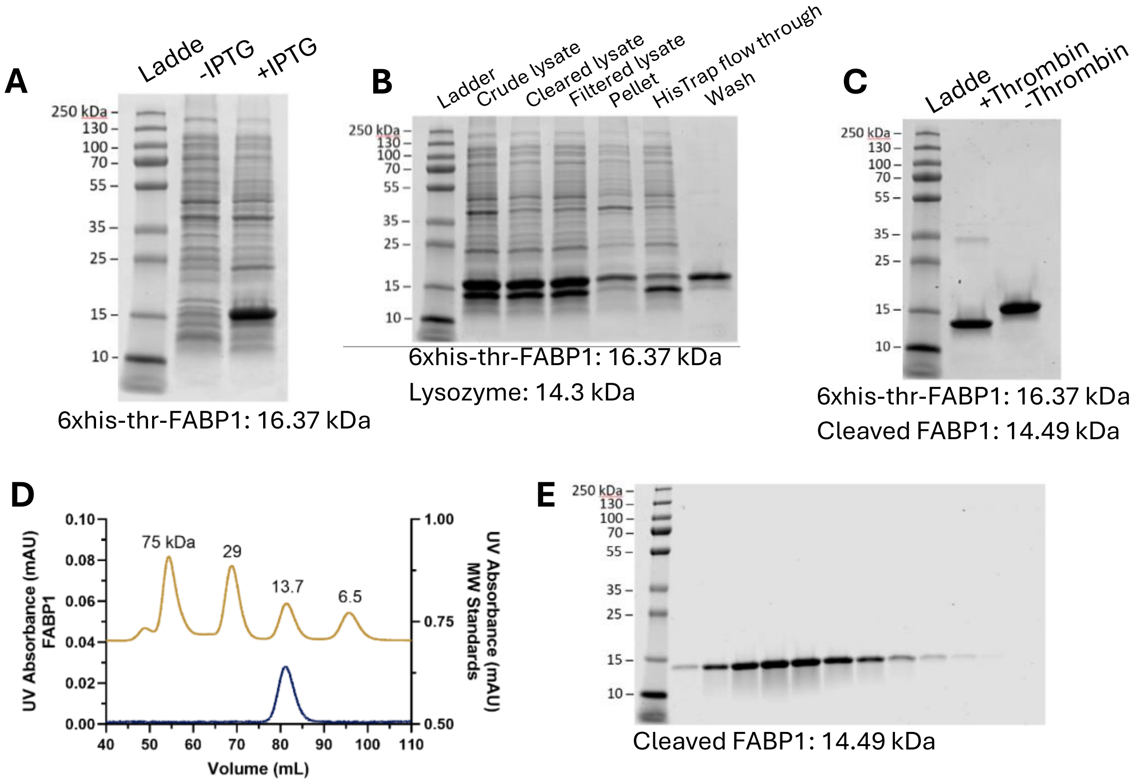


**Figure S2:** Expression and purification of FABP1. Coomassie stained SDS-PAGE gels of (**A**) whole cell *E. coli* lysate of his-tagged FABP1 induced with isopropyl β-D-1-thiogalactopyranoside (IPTG), (**B**) HisTrap purification steps and (**C**) thrombin cleavage of FABP1. Predicted molecular weights based on amino acid sequences are listed below the gel images. Panel (**D**) shows a gel filtration (Superdex 75) A280 chromatogram of monomeric thrombin-cleaved FABP1 post delipidation (dark blue trace) along with molecular weight standards (gold trace). The elution fractions (1 mL each) from the final gel filtration step are shown in the Coomassie stained gel in **E**.

Gel filtration and delipdation with Lipidex-5000 gravity flow column: Gel filtration was used to confirm monomeric protein while delipidation was done to ensure that there were no copurifying molecules and lipids present in the protein that could confound ligand binding analysis. In initial experiments, the his-tagged FABP1 was dialyzed overnight against 20 mM Tris pH 7.4 at 4°C, 500 mM NaCl to remove imidazole from the HisTrap elution step. The his-tagged FABP1 was then flash frozen and stored at -80°C until subsequent thrombin cleavage and delipidation steps. Cleavage of the his-tag was achieved by incubating his-tagged FABP1 with thrombin (0.03 U thrombin per 10 μg of FABP1) on ice for 18 hours. The cleaved protein (2 mL) was injected into a Superdex 75 size exclusion column (GE Healthcare, Chicago, IL) equilibrated with gel filtration buffer (20 mM Tris pH 7.4 at 4°C, 100 mM NaCl) using a DuoFlow FPLC at a flow rate of 0.5 mL/min. After injection, the flow rate was increased to 1 mL/min and FABP1 elution monitored by UV absorbance at 280 nm. 1 mL fractions were collected and the FABP1 containing fractions corresponding to the UV peak were verified by SDS-PAGE and Coomassie staining.

For delipidation of FABP1, 5 mL of Lipidex-5000 available as a methanol slurry (Perkin Elmer, Shelton, CT) was poured into a gravity flow column and washed with 10 column volumes of gel filtration buffer. The Lipidex-5000 was then conditioned in gel filtration buffer for at least 2 hours at 21°C. Gel filtration fractions containing FABP1 were pooled, passed 5 times through the Lipidex-5000 column and concentrated using an Amicon concentrator with a 10 kDa molecular weight cutoff (Sigma, St. Louis, MO). The protein was then flash frozen and stored in -80°C until native MS analysis. When this protein was analyzed using native MS several copurifying compounds were identified despite the delipidation protocol employed (Figure S3). This prompted us to further optimize the purification and delipidation protocol.

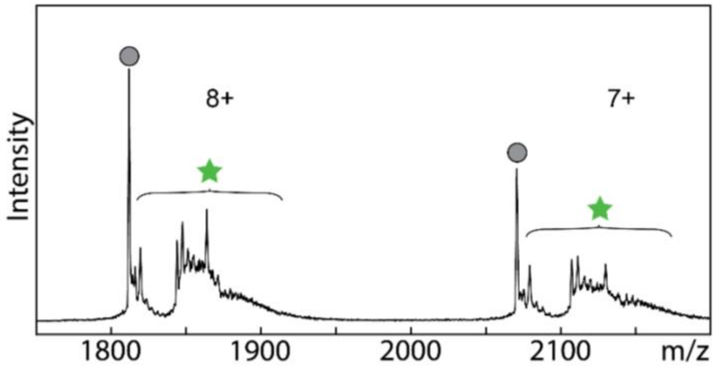


**Figure S3:** Native mass spectrum of FABP1 delipidated with 5 passes through a Lipidex-5000 gravity flow column. The 7+ and 8+ charge states for apo-FABP1 are denoted with circle markers. Star markers denote the region of the spectrum where *m/z*-shifted peaks are observed corresponding to co-purifying molecules.

Delipidation with Lipidex-5000 bead suspension: The effectiveness of delipidation using a Lipidex-5000 bead suspension method was tested based on previously published protocols (Wang *et al.*, 2017; Lai *et al.*, 2020). Lipidex-5000 is available as dry powder or as a slurry in methanol. The dry powder has been previously reported to effectively delipidate rat FABP2 and murine FABP4 (Wang *et al.*, 2017). To test whether the dry powder could be conditioned in-house to provide efficient delipidation of human FABP1, 1 g of Lipidex-5000 powder was resuspended in 4 mL of incubation buffer (10 mM potassium phosphate buffer at pH 7.4 with 150 mM KCl) and preconditioned by shaking for 2.5 hours at 37°C in a 15 mL Falcon tube. After preconditioning, his-tagged FABP1 dialyzed overnight into incubation buffer was added to the Lipidex-5000 suspension for a final volume of 10 mL and final FABP1 concentration of 0.8 mg/mL. The Lipidex-5000 slurry with FABP1 was incubated at 37°C with shaking for 2 hours. Visible precipitate was observed in the slurry after the 2 hour incubation. The slurry was then poured into gravity flow column, and the flow through was collected. The Lipidex-5000 was washed once with 5 mL incubation buffer and once with 5 mL 20 mM Tris pH 7.4 at 4°C with 100 mM NaCl. FABP1 concentrations in the slurry, flow through and washes were determined with A280 and recovery of FABP1 was assessed via SDS-PAGE and Coomassie staining. Analysis of the Coomassie stained gels showed that majority of the FABP1 remained in the Lipidex-5000 post washes suggesting the protein had precipitated and bound to the Lipidex-5000 during the 2 hour incubation

(Figures S4A). Despite extensive efforts, the FABP1 could not be eluted from the Lipidex-5000 slurry.

We then tested a shorter incubation time with Lipidex-5000 in a separate experiment. In this experiment, 1 mL of his-tagged FABP1 (3.8 mg/mL) in incubation buffer was added directly to 1 g of Lipidex-5000 (pre-conditioned as described above) in a 20 mL Poly-prep column (Bio-Rad, Hercules, CA). The FABP1 and Lipidex-5000 were agitated to mix and incubated at 21°C for 30 minutes. After 30 minutes, the flow through was collected and the Lipidex was washed once with 5 mL incubation buffer and once with 5 mL 20 mM Tris pH 7.4 at 4°C with 100 mM NaCl. Based on A280 and SDS-PAGE with Coomassie staining, the majority of the protein was recovered in the flow through or wash steps (Figure S4B). No precipitation of FABP1 was observed with the 30 minute incubation and a negligible amount lost to Lipidex-5000 binding. The extent of delipidation for the 30 minute incubation protocol was assessed with native protein MS (Figure 1B).

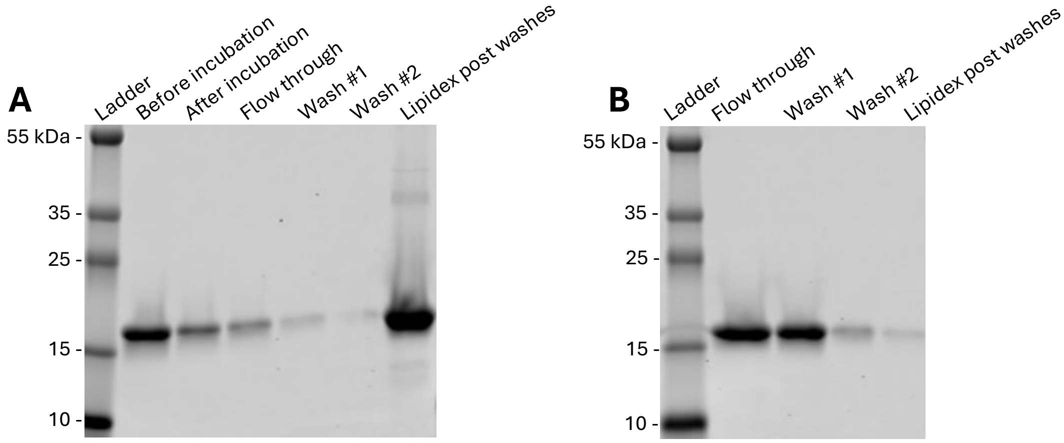


**Figure S4:** Stability of his-tagged FABP1 with Lipidex-5000. Coomassie stained SDS-PAGE gels of nickel purified his-tagged FABP1 incubated with Lipidex-5000 slurry prepared from powder for (**A**) 2 hours at 37°C and (**B**) 30 minutes at 21°C. Visible precipitate was observed with the 2 hour incubation and the majority of FABP1 was bound to Lipidex-5000 (A, last lane). In contrast, no precipitate was observed with the 30 minute incubation and the majority of FABP1 was recovered in the flow through and washes (B, lane 2 and 3).

<ins>Delipidation with Butanol:</ins> Based on the preliminary experiments, Lipidex-5000 was not fully effective in removing all copurifying compounds from hFABP1 and hence more extensive delipidation protocols with treatments with 1:1 (v/v) butanol or 1:3 (v/v) butanol were tested. His-tagged FABP1 dialyzed into incubation buffer (10 mM potassium phosphate buffer at pH 7.4 with 150 mM KCl) was used to test butanol treatments. 300 µL (1:3) or 1 mL (1:1) of butanol were added to 1 mL of his-tagged FABP1 (3.8 mg/mL) in incubation buffer in a glass tube and the mixture was rocked for 10 minutes at 21°C. After 10 minutes, the solution was centrifuged at low speed for 30 seconds to separate the organic and aqueous phases. The organic phase was discarded and butanol treatment was repeated another 2 times for the 1:1 butanol treatment and additional 3 times for the 1:3 butanol treatment. The number of extractions was decided based on the decrease in aqueous phase volume after butanol treatment due to water partitioning to the butanol. The final

volume of the aqueous phase post butanol treatment was approximately half of the initial volume and minimal protein was lost based on A280 measurements. The extent of delipidation for the his-tagged FABP1 was then verified via native protein MS (Figure 1). The least amount of co-purifying molecules (CPMs) was observed in the MS spectrum of FABP1 treated with 1:1 butanol.

Delipidation with a combination of Lipidex-5000 and butanol: Since residual CPMs were still bound to FABP1 based on the native mass spectra of individual treatments with Lipidex-5000 or butanol (Figure 1), a combination of sequential Lipidex-5000 and butanol treatments was tested. Swelling of Lipidex-5000 was better achieved with sonication in 20% methanol and longer conditioning times compared to the conditioning protocol described above. Due to these technical considerations, Lipidex-5000 available as a methanol slurry was chosen for delipidation and the following conditioning protocol was used. 2 g of Lipidex-5000 in methanol was transferred to a 50 mL Falcon tube, pelleted at 500 g for 5 minutes and the methanol decanted. 20 mL of gel filtration buffer with 20% methanol was added to the Lipidex-5000 and the Lipidex-5000 was resuspended by stirring with a glass Pasteur pipette. The Lipidex-5000 was then incubated in a bath sonicator for 10 minutes. After sonication, the Lipidex-5000 was pelleted at 500 g for 5 minutes, the buffer with methanol was removed and the Lipidex-5000 washed with 20 mL of gel filtration buffer twice before resuspending to a final volume of 20 mL of gel filtration buffer (0.1 g Lipidex-5000 per mL of buffer). The Lipidex-5000 was resuspended in buffer by stirring with a glass Pasteur pipette, sonicated for an additional 10 minutes then incubated in buffer overnight at 21° C. The overnight conditioned Lipidex-5000 was poured into a 10 mL Poly-prep column (0.1 g Lipidex-5000 per 1 mg FABP1) and the buffer removed by gravity flow before adding FABP1 for delipidation. The FABP1 was incubated with the Lipidex-5000 with rocking at 21°C for 30 minutes and then eluted from the column via gravity flow.

To test different combinations of sequential Lipidex-5000 and butanol treatments, purified FABP1 post thrombin cleavage and gel filtration (as described above) was delipidated with either butanol (1:1) for 3 rounds or incubation for 30 minutes with Lipidex-5000 (as described above). For butanol extracted FABP1, subsequent 30 minute incubations with Lipidex-5000 were performed for 3 rounds. For FABP1 initially treated with Lipidex-5000, 3 sequential rounds of butanol extraction were performed. Samples of FABP1 were collected after each treatment round and delipidation was assessed using native protein MS. The combination of butanol extraction followed by Lipidex-5000 treatment showed the least amount of CPMs based on MS spectra (Figure S5). Subsequent rounds of Lipidex did not further improve delipidation after the first Lipidex treatment (data not shown). The final protocol chosen for delipidation was extraction for 3 times with 1:1 butanol followed by a 30 minute incubation with Lipidex-5000. The final protocol is described below.

Final purification and delipidation protocol: In the final protocol (Figure 2), the peak fractions from the HisTrap purification were combined and diluted with an equivalent volume of 20 mM Tris pH 7.4 at 4°C, 100 mM NaCl. The concentration of FABP1 was measured using a Nanodrop A280. The N-terminal his-tag was then cleaved by adding 0.03 U of thrombin protease per 1 µg of FABP1 and incubating the mixture in a water bath for 1 hour at 37°C. Complete cleavage of the N-terminal tag was confirmed via SDS-PAGE and Coomassie staining (Figure

S2C) and with an anti-His (mouse anti-His antibody from Qiagen, Valencia, CA) western blot. The cleaved protein (2 mL) was then injected into a Superdex 75 size exclusion column (GE Healthcare, Chicago, IL) equilibrated with gel filtration buffer (10 mM potassium phosphate pH 7.4, 150 mM KCl) using a DuoFlow FPLC at a flow rate of 0.5 mL/min. After injection, the flow rate was increased to 1 mL/min and FABP1 elution monitored by UV absorbance at 280 nm. 1 mL fractions were collected and the FABP1 containing fractions corresponding to the UV peak were verified by SDS-PAGE and Coomassie staining (Figure S2D and S2E). Monomeric FABP1 was verified using gel filtration and by comparison of the molecular weight to standards from a calibration kit (Cytiva, Marlbough, MA). FABP1 containing fractions (up to 4 mL) were pooled for removal of copurifying lipids using the final delipidation protocol.

For delipidation an equivalent volume of butanol was added to the FABP1 containing fractions. The butanol and FABP1 mixture was ro cked at room temperature in a glass tube for 10

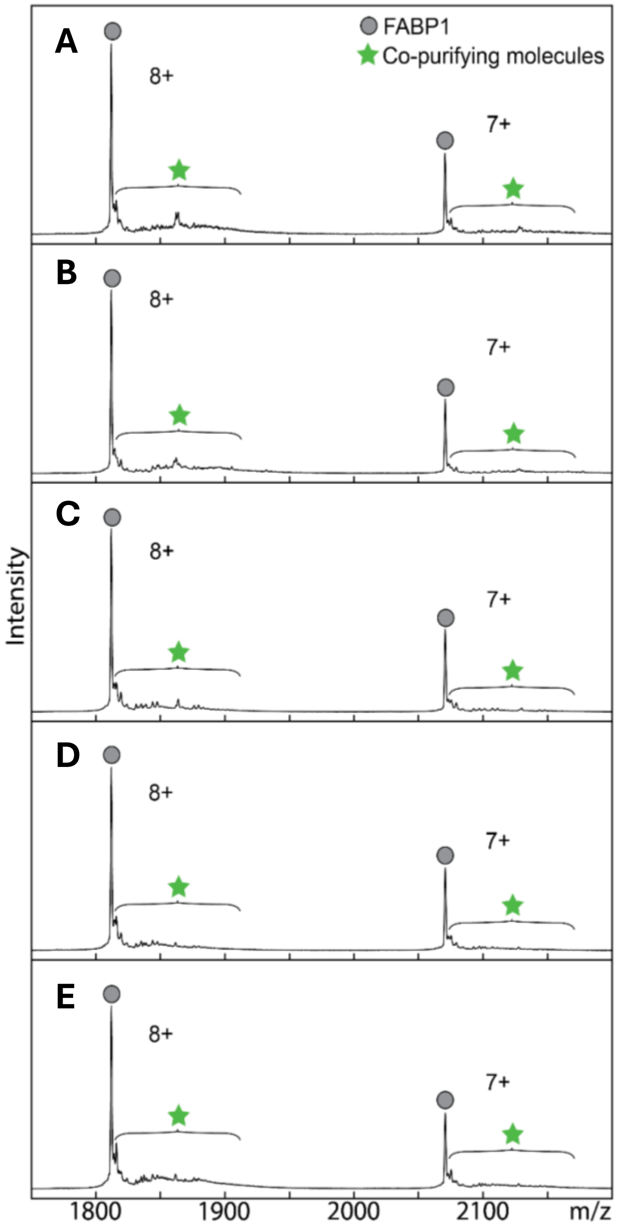


**Figure S5:** Comparison of FABP1 delipidation with sequential treatments of Lipidex-5000 and butanol. Native mass spectra are shown for FABP1 post thrombin cleavage and gel filtration (**A**), then subsequent treatment with 3 rounds of 1:1 (v/v butanol to FABP1 solution) butanol (**B**), 30 minute incubation with Lipidex-5000 (**C**), a combination of 3 rounds 1:1 butanol then 30 minute incubation with Lipidex-5000 (**D**), or a combination of 3 rounds of 30 minute incubation with Lipidex-5000 then 3 rounds of 1:1 butanol (**E**).

minutes. The mixture was then centrifuged at low speed for 30 seconds to separate butanol and aqueous phases and the butanol phase was removed with a Pasteur pipette. Butanol extraction was repeated for a total of 3 rounds. The aqueous phase containing FABP1 was then transferred to a 10 mL Poly-Prep chromatography column (Bio-Rad, Hercules, CA) containing Lipidex-5000 (0.1 g of Lipidex-5000 per 1 mg of FABP1) pre-conditioned with sonication and overnight incubation as described above. FABP1 was incubated with Lipdex-5000 with rocking at 21°C for 30 minutes. After incubation with Lipidex-5000, traces of butanol were removed by gel filtration chromatography. The Superdex 75 size exclusion column was equilibrated with 10 mM potassium phosphate pH 7.4 with 150 mM KCl. FABP1 containing gel filtration fractions were pooled, stored on ice and the concentration of delipidated FABP1 was quantified via bicinchoninic acid (BCA) assay prior to adding 0.5 mM DTT.

### Initial Characterization of DAUDA Binding to FABP1

Solutions of DAUDA and FABP1 were prepared in 2 mL volumes. Stock solutions of DAUDA were prepared in methanol and DAUDA concentrations confirmed using a Cary 60 UV-Vis spectrophotometer (Agilent, Santa Clara, CA) assuming a DAUDA molar absorption coefficient of 4400 $M^{-1}$ $cm^{-1}$ at 335 nm in methanol (Thumser *et al.*, 1996). Fluorescence spectra were collected using a Cary Eclipse fluorescence spectrophotometer (Agilent, Santa Clara, CA) and fluorescence measurements were taken in a 4 mL clear quartz cuvette. The scan rate was set to medium (600 nm/min) and the photomultiplier tube voltage was set to high (800 V). The fluorescence spectrum of DAUDA was initially verified in the presence and absence of FABP1. Briefly, solutions of DAUDA (0.08, 0.28, 0.68 µM) in the absence or presence of FABP1 (5 µM) were made in 50 mM potassium phosphate buffer, pH 7.4 with 100 mM KCl. DAUDA binding to FABP1 was monitored via the enhancement of fluorescence in the presence of FABP1 using an excitation wavelength of 335 nm and emission was monitored from 400-700 nm (Figure S6).

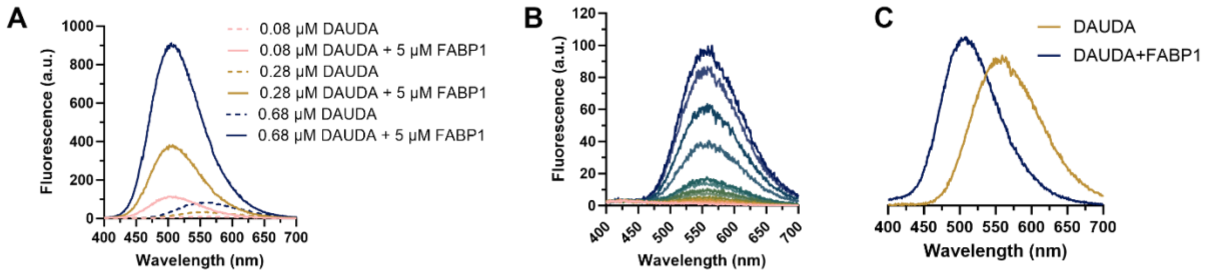


**Figure S6:** Fluorescence emission spectra and basis spectra of DAUDA in the presence and absence of FABP1. (**A**) Fluorescence spectra for different concentrations of DAUDA alone (0.08, 0.28, 0.68 µM) in buffer (dotted lines) and in the presence of 5 µM FABP1 (solid lines). (**B**) Fluorescence spectra of free DAUDA in buffer. Each colored line represents a different concentration of DAUDA (0.02-1.03 µM). (**C**) Basis spectra used for spectral deconvolution of fluorescence spectra. Gold spectrum is the basis spectra for DAUDA in solution and dark blue spectrum is the basis spectrum of DAUDA-FABP1.

**Analysis of Titration Spectra by Singular Value Decomposition**

Singular value decomposition (SVD) is an analytical technique based on linear algebra that can be used to deconvolute fluorescence titration spectra from equilibrium binding experiments. A general approachable review of the linear algebra, theory and application of SVD can be found in (Hendler and Shrager, 1994). Spectral filtering and deconvolution of fluorescence spectra from titration experiments with DAUDA was done with a previously described in house SVD program (Nath *et al.*, 2008). The program and source code are available at the following web address: http://marvin.mchem.washington.edu/pca/. Titration emission spectra from a single experiment were formatted to tab-delimited text files (.txt) where the first column indicated the wavelength and each subsequent column were the observed fluorescence values corresponding to a single concentration of the ligand being titrated. An example of properly formatted data can be found at the web address above. Each experiment was initially analyzed using the default number of components (4). The number of components chosen to be retained in the final analysis was based on inspection of the spectral components and the singular values (scree plot) that were above baseline values (i.e noise). Explicit basis spectra of DAUDA alone in solution and DAUDA-FABP1 (Figure S7C) were uploaded into the program for use in deconvolution of titration spectra. A flat line was included in the basis spectra file to account for any residual fluorescence that was not described by the basis spectra of DAUDA or DAUDA-FABP1. The specific fluorescence for DAUDA alone and DAUDA-FABP1 were obtained from the matrix "Concentration vectors of basis spectra." These values were plotted against the concentrations of titrated ligand to generate binding curves for reverse and forward titrations and DAUDA displacement titration experiments.

Construction of Basis Spectra: The basis spectra for DAUDA alone in solution and DAUDA-FABP1 were obtained from experimental spectra of titrations with DAUDA alone or DAUDA in the presence of excess FABP1. The experimental titration spectra were first filtered through the in house SVD program. For the basis spectrum of DAUDA alone, the data weight values of the first two components found in the matrix "Concentration vectors of spectral components" were obtained corresponding to the concentration of DAUDA at 1.03 µM in assay buffer (Figure S7B). These data weight values were then used as the coefficients in the option "choose to estimate basis spectra based on calculated spectral components" to reconstruct the basis spectrum. The basis spectrum generated under "Basis spectra used for deconvolution" was normalized to the peak area and used as explicit basis spectrum for SVD analysis (Figure S7C). The basis spectrum of DAUDA bound with FABP1 (DAUDA-FABP1) was similarly constructed. Fluorescence spectrum where all DAUDA is expected to be bound to FABP1 was chosen from the reverse titration (0.05 µM DAUDA with 1.5 µM FABP1) and forward titration (0.02 µM DAUDA with 0.3 µM FABP1) (Figure 3) to generate the basis spectrum of DAUDA-FABP1. The basis spectra of the unique components of (R)- and (S)-flurbiprofen (Figure S8) could not be isolated experimentally and were constructed based on visual inspection of the filtered spectral components.

### Determination of *Scale* for COPASI Model Fitting of Arachidonic Acid and Drug Titration Data

The specific fluorescence of DAUDA-FABP1 from arachidonic acid (AA) and drug titration experiments determined by SVD was normalized using the following equation:

$$F_{norm} = \frac{F_{[Ligand]}}{F_0} \times 100 \quad \text{(S1)}$$

In equation S1, $F_{[Ligand]}$ is the specific fluorescence of DAUDA-FABP1 at a given ligand concentration and $F_0$ is the specific fluorescence of DAUDA-FABP1 in the absence of ligand and with 0.5 and 0.3 µM total DAUDA and FABP1, respectively. Therefore, the maximum normalized fluorescence in the absence of ligand was 100 arbitrary fluorescence units. The maximum normalized fluorescence in the absence of ligand could then be described by the following equation:

$$100 = Scale \cdot ([FABP1 - DAUDA] + 2 \cdot [FABP1 - DAUDA - DAUDA]) \quad \text{(S2)}$$

Equation S2 was then rearranged to:

$$Scale = \frac{100}{[FABP1 - DAUDA] + 2 \cdot [FABP1 - DAUDA - DAUDA]} \quad \text{(S3)}$$

The concentrations of [FABP1-DAUDA] and [FABP1-DAUDA-DAUDA] were determined with COPASI using the two-site sequential binding model for DAUDA. At equilibrium, the predicted concentrations of [FABP1-DAUDA] and [FABP1-DAUDA-DAUDA] were 0.179 and 0.005 µM, respectively. *Scale* was then determined using equation S3, yielding a value of 529.

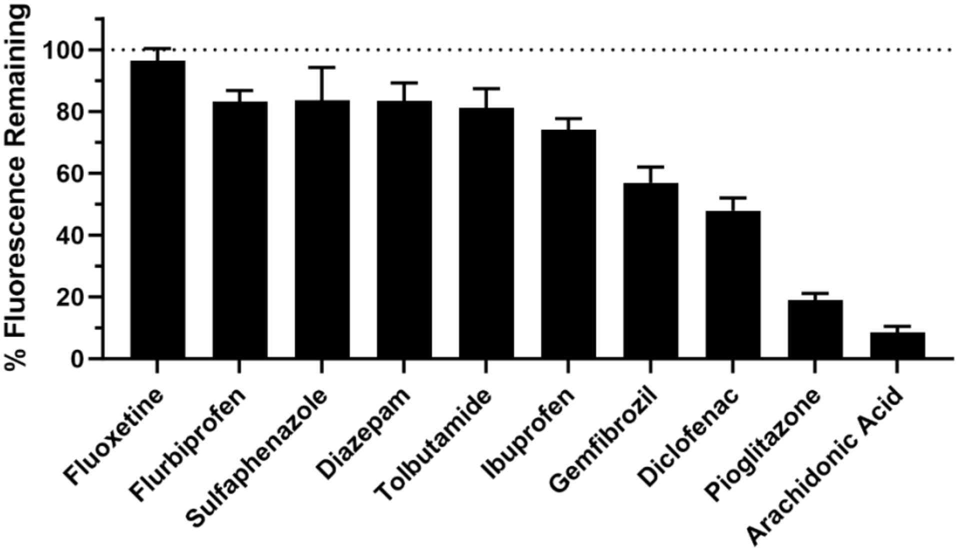


**Figure S7:** Screening of DAUDA displacement by potential drug ligands. Each drug was screened at 30 µM concentration added to FABP1 (0.3 µM) prebound with DAUDA (0.5 µM). The % fluorescence remaining based on the decrease in fluorescence intensity at 500 nm relative to a no ligand control is shown (mean ± standard deviation) from replicate experiments performed on 3 separate days.

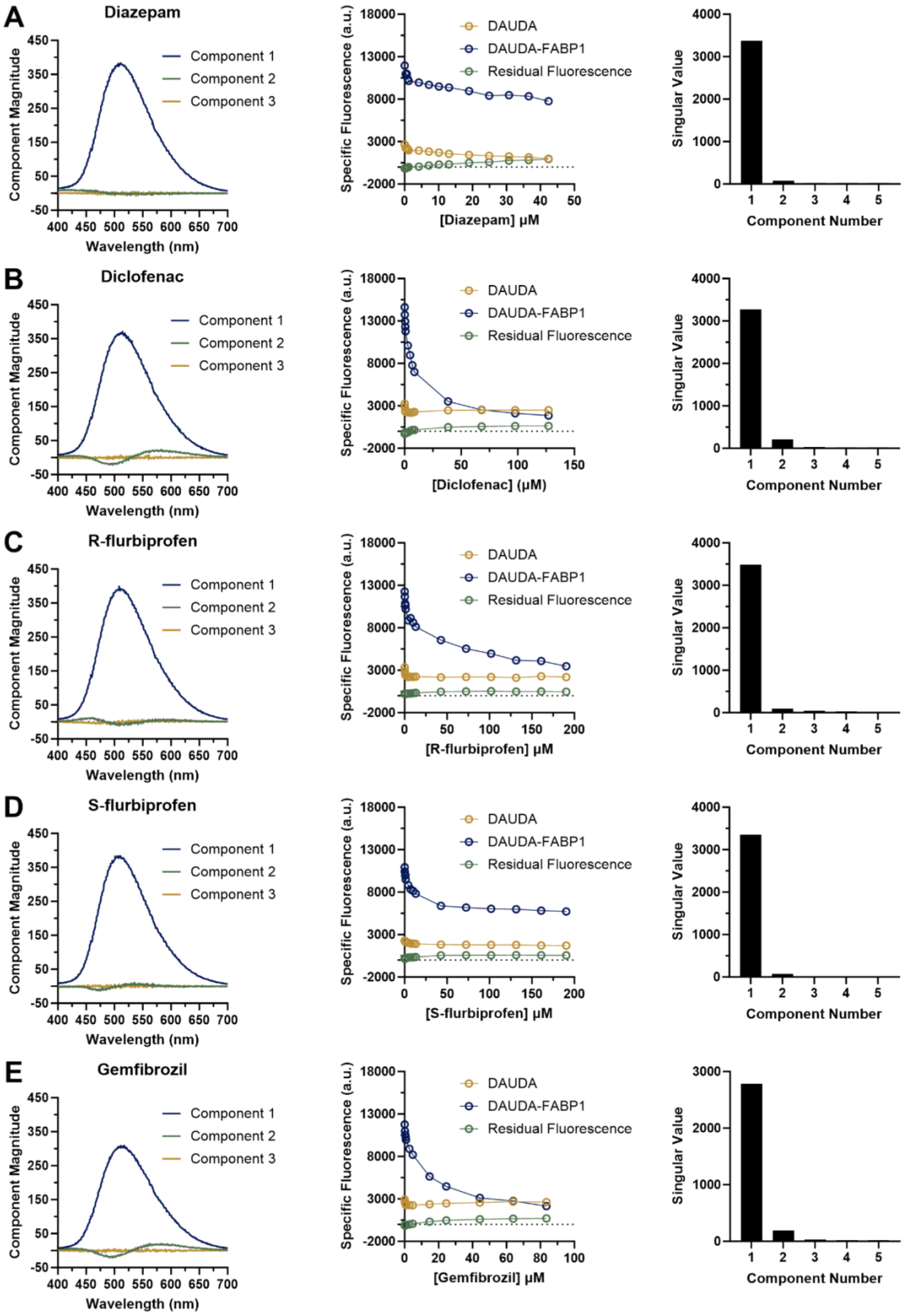
A
Diazepam
B
Diclofenac
C
R-flurbiprofen
D
S-flurbiprofen
E
Gemfibrozil
Component Magnitude
Wavelength (nm)
Component 1
Component 2
Component 3
Specific Fluorescence (a.u.)
DAUDA
DAUDA-FABP1
Residual Fluorescence
[Diazepam] μM
[Diclofenac] (μM)
[R-flurbiprofen] μM
[S-flurbiprofen] μM
[Gemfibrozil] μM
Singular Value
Component Number

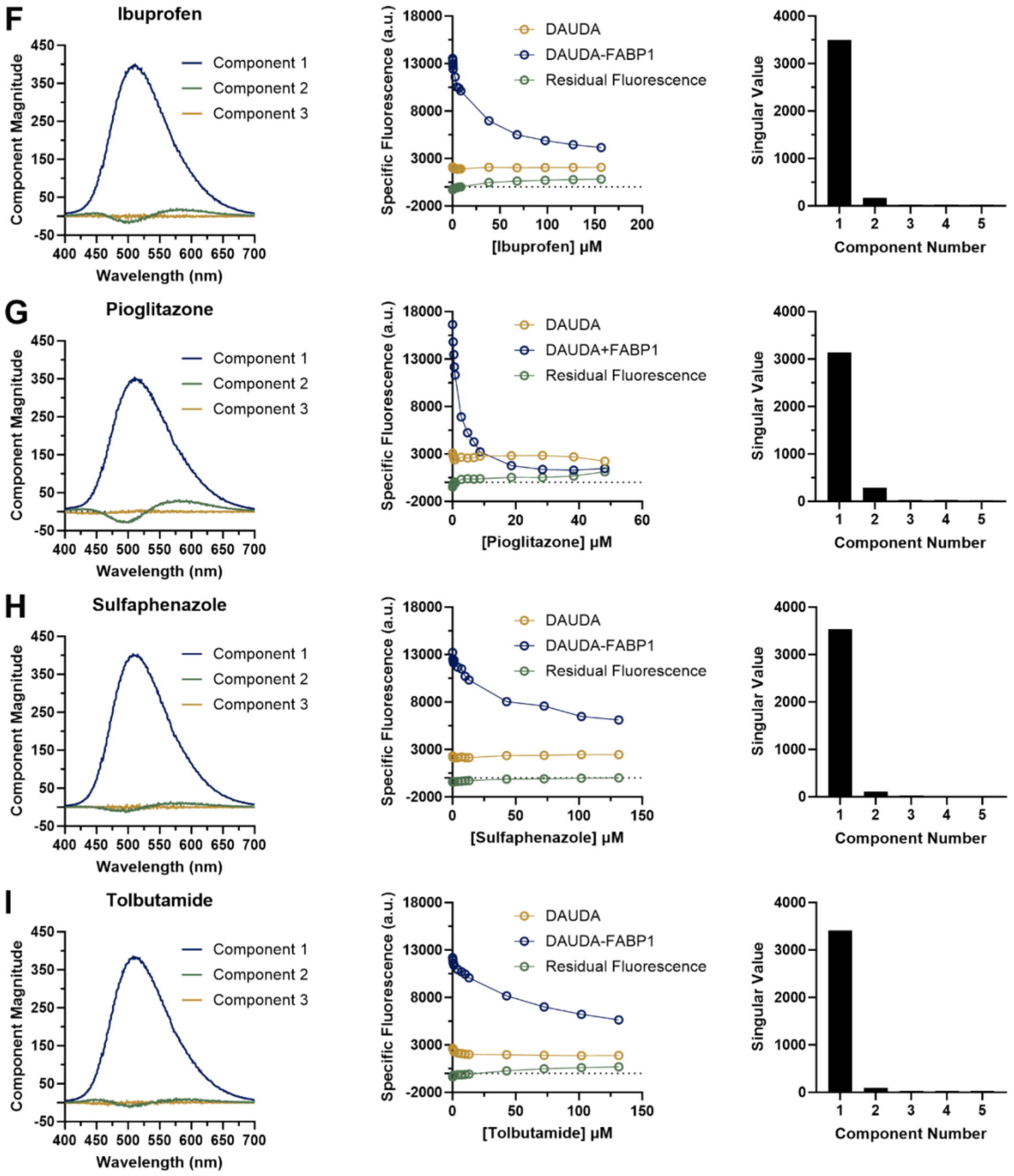


**Figure S8:** SVD analysis of DAUDA displacement titration for all drug ligands tested. Each lettered panel (A-I) shows the SVD analysis for a representative replicate titration for each drug ligand tested in DAUDA displacement assays. For each drug titration, plots are shown for (left) the first three spectral components identified from SVD, (middle) change in specific fluorescence of DAUDA and DAUDA-FABP1 with increasing ligand concentration and (right) a scree plot with the strength (singular value) of the first 5 components from SVD. The primary component (blue) for all ligands shown match the emission peak of DAUDA-FABP1 ($\lambda_{em}$ 509 nm).

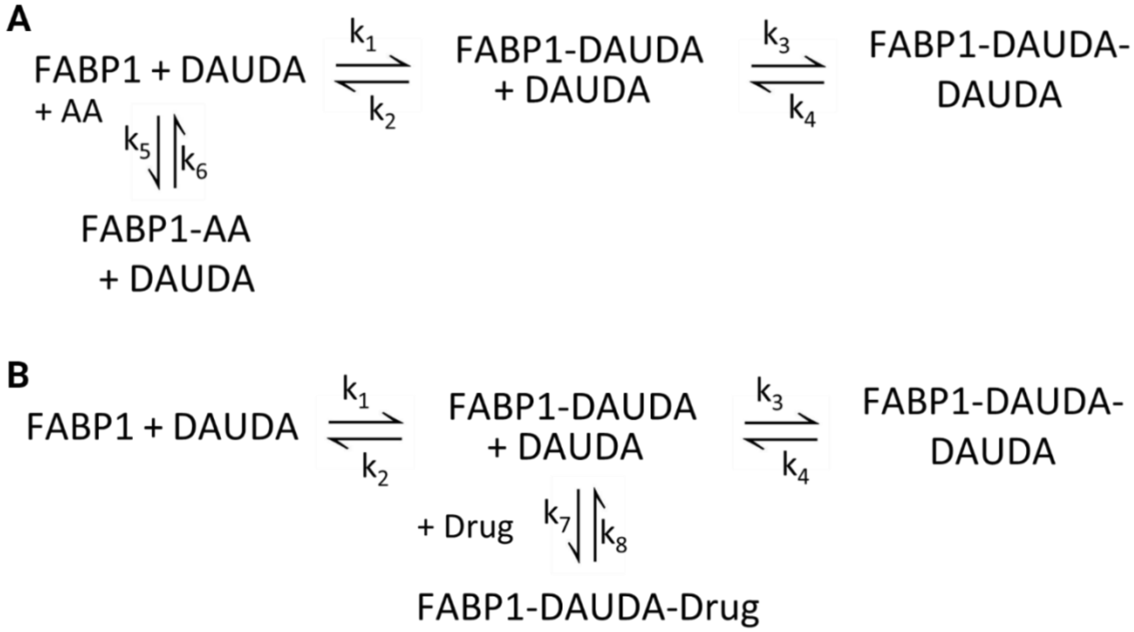


**Figure S9:** Kinetic schemes for arachidonic acid (AA) and drug binding models used in COPASI. (**A**) Competitive binding model for AA. AA competes with DAUDA for binding to FABP1 occupying both DAUDA binding sites. (**B**) Ternary binding model for drugs. Drugs bind at a second binding site in FABP1 without displacing DAUDA to form a ternary complex. Schemes created by BioRender.com.

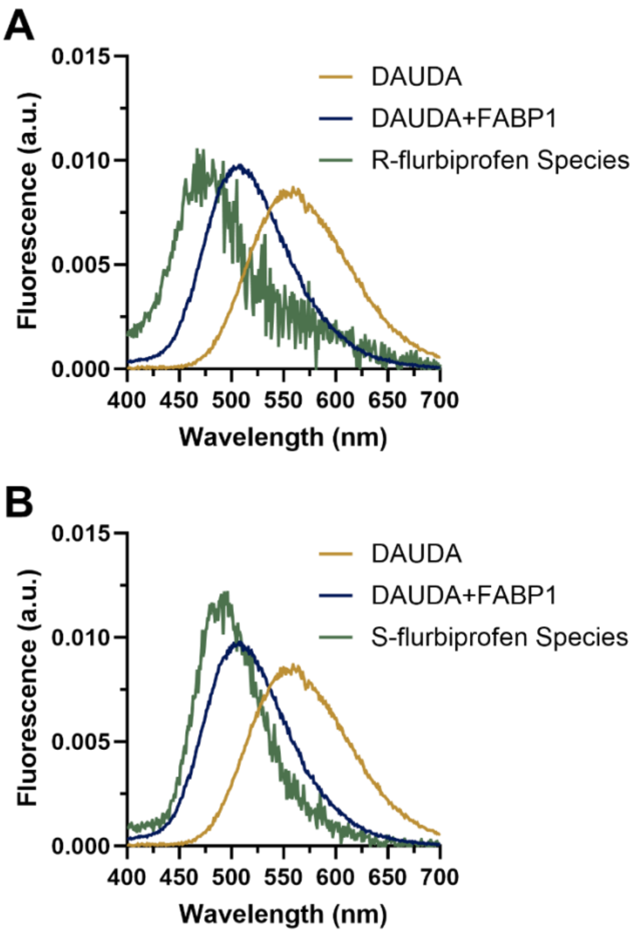


**Figure S10:** (R)- and (S)-flurbiprofen basis spectra. Basis spectra for (**A**) (R)- and (**B**) (S)-flurbiprofen used for spectral deconvolution of titration data. DAUDA (gold) and DAUDA-FABP1 (blue) basis spectra were experimentally determined as described above. The basis spectra for bound (R)- and (S)-flurbiprofen (green) were generated based on unique spectral components identified from SVD analysis of titration spectra with (R)- and (S)-flurbiprofen.

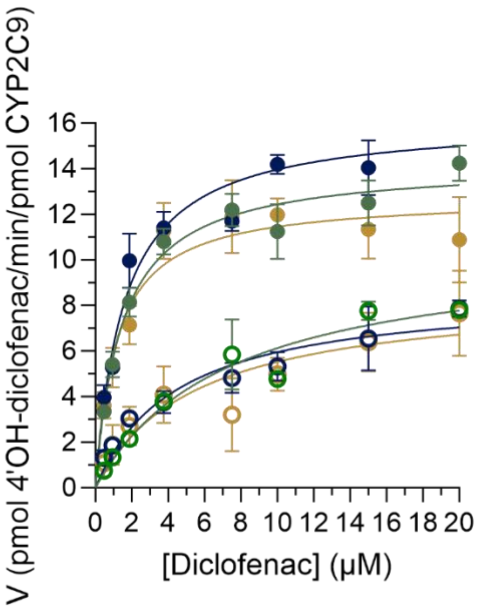


**Figure S11:** Formation of 4’OH-diclofenac from diclofenac by CYP2C9 in the presence (open circles) and absence (solid circles) of 20 µM FABP1 is shown. Paired replicate experiments from three separate days are shown in dark blue, green and gold. The x-axis shows the nominal concentration of diclofenac added to the incubations.
